# CTCF and cohesin promote focal detachment of DNA from the nuclear lamina

**DOI:** 10.1101/2021.09.13.460079

**Authors:** Tom van Schaik, Ning Qing Liu, Stefano G. Manzo, Daan Peric-Hupkes, Elzo de Wit, Bas van Steensel

## Abstract

Lamina associated domains (LADs) are large genomic regions that are positioned at the nuclear lamina (NL). It has remained largely unclear what drives the positioning and demarcation of LADs. Because the insulator protein CTCF is enriched at LAD borders, it was postulated that CTCF binding could position a subset of LAD boundaries, possibly through its function in stalling cohesin and hence preventing cohesin to invade into the LAD. To test this, we mapped genome – NL interactions in mouse embryonic stem cells after rapid depletion of CTCF and other perturbations of cohesin dynamics. CTCF and cohesin contribute to a sharp transition in NL interactions at LAD borders, whilst LADs are maintained after depletion of these proteins, also at borders marked by CTCF. CTCF and cohesin may thus reinforce LAD borders, but do not position these. CTCF binding sites within LADs are locally detached from the NL and enriched for accessible DNA and active histone modifications. Remarkably, even though NL positioning is strongly correlated with genome inactivity, this DNA remains accessible after the local detachment is lost following CTCF depletion. At a chromosomal scale, cohesin depletion and cohesin stabilization (depletion of the unloading factor WAPL) quantitatively affect NL interactions, indicative of perturbed chromosomal positioning in the nucleus. Finally, while H3K27me3 is locally enriched at CTCF-marked LAD borders, we find no evidence for an interplay between CTCF and H3K27me3 on NL interactions. Combined, these findings illustrate that CTCF and cohesin do not shape LAD patterns. Rather, these proteins mediate fine-tuning of NL interactions.

## Background

The nuclear lamina (NL) is positioned near the inner nuclear membrane and consists of a protein meshwork of A- and B-type lamins and a variety of other proteins. Peripherally positioned chromatin interacts with the NL and comprises approximately a thousand large genomic domains that are up to 10 Mb in size [1] [1]. These lamina-associated domains (LADs) are strongly depleted for active genes and are thought to provide a backbone for genome organization. LADs are highly conserved between cell types and species, but the mechanisms that underlie genome positioning at the NL remain poorly understood [reviewed in 2, 3, 4].

LADs are strongly enriched for features of heterochromatin. Besides having a low gene density and typically low gene expression levels, LADs replicate late in S-phase, are visible by electron microscopy as densely packed chromatin and are enriched for the histone modifications H3K9me2 and H3K9me3 [1, 5–9]. These heterochromatin characteristics are important because gene activation, chromatin decondensation and H3K9me2/3 perturbation all result in a loss of DNA – NL interactions [7, 10–14]. LAD interactions with the NL are mediated by various proteins in a redundant manner, including but probably not limited to Lamin B receptor (LBR), Lamin A and CEC-4 (the latter only in *C. elegans*) [13, 15]. Differences in NL interactions between cell types are likely caused by variable epigenetic landscapes and protein compositions of the NL.

One particular aspect of LADs that remains poorly understood is the sharp transition in NL interactions at LAD borders, which suggests the existence of specific mechanisms to define these borders. LAD borders are enriched for active promoters, CpG islands (often at active promoters) and binding of the insulator protein CTCF [1]. While there is increasing evidence that gene activity results in local detachment from the NL [11, 14], which could explain a sharp transition at some borders, the role of CTCF on LAD border positioning is still unclear. Because few borders are marked by both CTCF and active promoters, it was postulated that CTCF may independently demarcate at least a subset of LAD borders [1, 12]. In support of this hypothesis, a deletion of a ∼30 kb LAD border region at the *Tcrb* locus resulted in decreased NL interactions up to 100 kb inside the LAD, and this behavior was almost completely recapitulated by deletion of a ∼4.5 kb region that included all three CTCF binding sites at this border [16]. However, a genome-wide view of a possible causal role of CTCF in LAD border formation is still lacking.

CTCF is a broadly expressed and essential zinc-finger protein that interacts with the ring-shaped cohesin complex on the genome [17, 18]. Once loaded on DNA, cohesin is thought to establish long-range DNA contacts through a processive increase in chromatin loop size, a process known as extrusion [19, 20, reviewed in 21]. Extrusion is halted by the interaction of the SA1/2-RAD21 interface of the cohesin complex with the N-terminal regions of CTCF [22, 23]. Cohesin is actively loaded by the NIPBL/MAU2 complex and unloaded by the cohesin release factor WAPL, to maintain a continuous cycle of loop extrusion [24, 25], which is required to preserve distal gene regulation [26]. Remarkably, besides having extended DNA loops, WAPL-knockout cells also have somewhat fragmented LADs [25]. However, the mechanism by which WAPL loss affects NL interactions remains to be elucidated.

In addition to the genomic features discussed above, in some cell types H3K27me3 is also enriched near LAD borders [1, 12]. H3K27me3 and CTCF could be involved in LAD demarcation together, given that both were reported to be involved in peripheral positioning of ectopically integrated LAD fragments [12]. However, the role of H3K27me3 may be cell-type specific, as this mark is not correlated with NL interactions in all cell types [10, 27]. Additionally, H3K27me3 is associated with dynamic LADs during the cell cycle [28], oncogene-induced senescence [29] and strain-induced genome repositioning [30].

Two recent developments now allowed us to investigate the interplay between LAD borders and CTCF and cohesin. First, fusion of endogenous proteins with auxin-inducible degrons (AID) tags mediate rapid and near-complete protein depletion, which is particularly useful for essential proteins or complexes such as CTCF and cohesin [31, 32]. Second, application of proteinA-DamID (pA-DamID) results in mapping of genome-NL interactions with high temporal resolution, which is required to study the effects of rapid protein depletion [33]. Here, using a combination of these two tools in mouse embryonic stem cells (mESCs), we show that CTCF and cohesin mediate local detachment from the NL, but have limited effects on LAD border positioning. On a chromosomal scale, perturbation of cohesin dynamics induces quantitative changes of NL interactions, which signifies that cohesin affects radial positioning of the genome. Finally, pharmacological depletion of H3K27me3 does not affect NL interactions in mESCs, even in combination with CTCF depletion.

## Results

### LAD borders are enriched for CTCF binding in multiple cell lines

We first set out to explore the correlation between CTCF binding (as detected by ChIP-seq) and LAD borders in more detail by comparing these in two mouse and four human cell lines with available NL interaction maps (obtained by DamID). CTCF binding is consistently depleted within LADs and locally enriched near LAD borders, peaking 10kb outside LADs (Fig 1A). CTCF is enriched at cell-type specific borders and at LAD borders shared between cell types (Fig S1A), although shared borders often have overall slightly elevated CTCF density. Thus, CTCF enrichment at LAD borders is not specific to either facultative or constitutive LADs.

**Figure 1.**
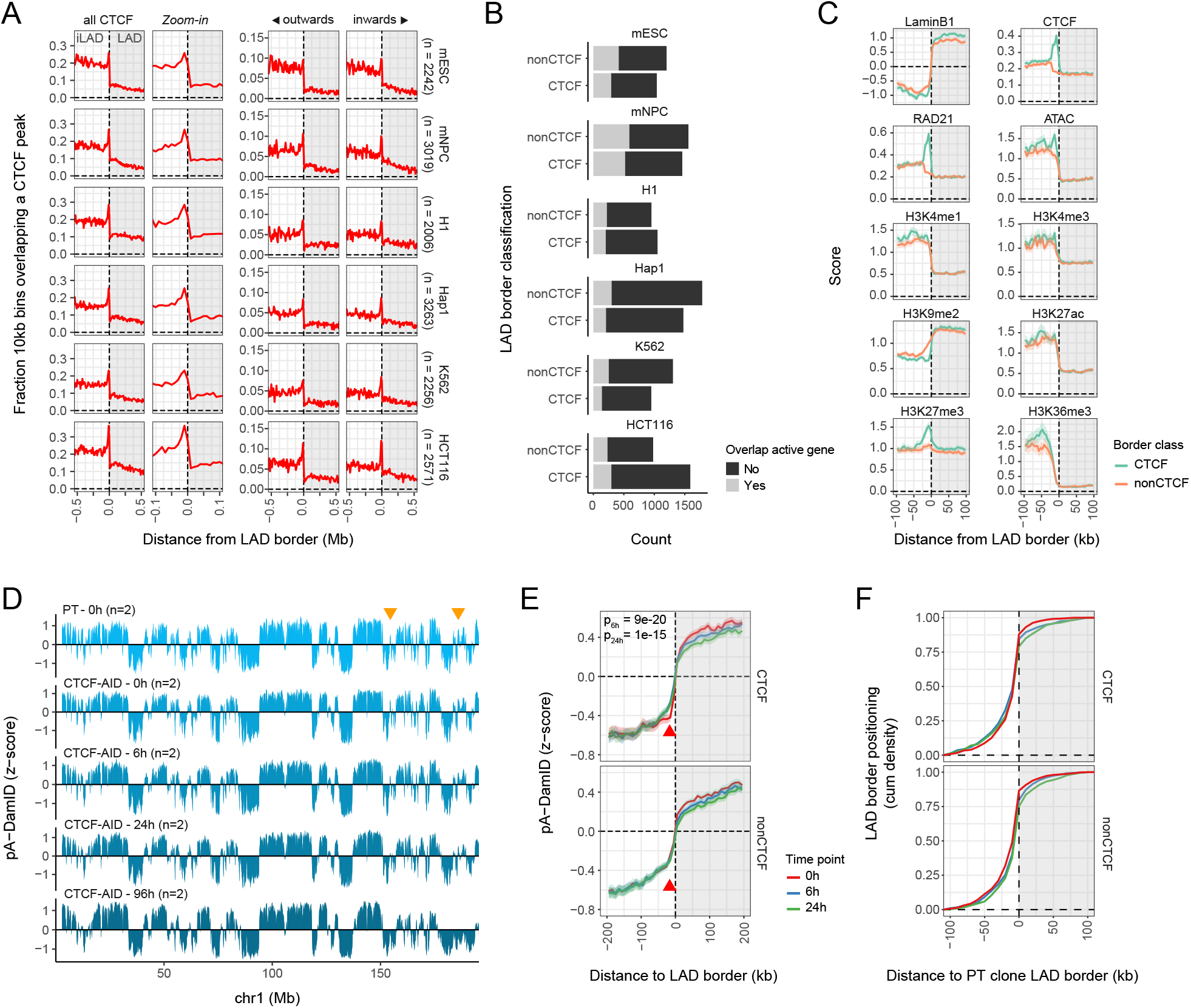
NL interactions are largely maintained upon CTCF depletion. **(A)** Positioning of CTCF binding sites around LAD borders in various mouse and human cell lines. Data are from refs [25, 26, 33, 66, 67]. LAD borders are defined based on DamID data binned at 10 kb bins, where “n” shows the number of LAD borders for every cell type (see Methods for more details). CTCF motif orientation at CTCF binding sites was used to assign binding directionality relative to the LAD. **(B)** Classification of LAD borders in CTCF and non-CTCF groups. LAD borders are classified as CTCF borders if a CTCF site is positioned within 20kb outside of the LAD (overlapping with the enriched CTCF density). LAD borders close to active genes (within 10kb, irrespective of their orientation) were removed from downstream analyses to prevent confounding effects. Active genes are defined as genes with an FPKM score higher than 1. Data are from refs [25, 26, 37, 67–70]. **(C)** Average scores for LaminB1 DamID (log_2_-ratio over Dam), CTCF and cohesin binding (ChIP-seq coverage), and various publicly available epigenetic data sets around mESC LAD borders, classified by CTCF presence. Data are from [26, 71–75]. Solid lines and shaded areas represent the mean signals and 95%-confidence intervals of the mean, respectively. **(D)** LaminB1 pA-DamID z-scores along a representative chromosome for mESC parental clone (PT) and for a time course of IAA addition in CTCF-AID cells (0, 6, 24h and 96 hours of CTCF depletion). Sequencing reads are counted and normalized in 10 kb bins. Data tracks are averages of *n* biological replicates. Orange arrows highlight example regions with variable signal following CTCF depletion. **(E)** Average LaminB1 pA-DamID z-scores of the CTCF-AID samples described in (D) (except 96h) around LAD borders (from mESC PT pA-DamID data, Fig S2E). The solid line and the shaded area represent the mean signal and 95%-confidence interval of the mean, respectively. The red arrow highlights the position of the CTCF enrichment that shows a local increase in pA-DamID z-scores after addition of IAA. The changes in NL interactions at the red arrow were calculated for all LAD borders, after which a Wilcoxon test was used to test for statistical significance between borders with and without CTCF. The Benjamini-Hochberg multiple testing correction was applied afterwards. **(F)** Cumulative density profile of LAD border positioning, as determined by hidden Markov modeling of the CTCF-AID samples described in (D) (except 96h), relative to the nearest LAD border from the PT clone. A distance cutoff of 100 kb was used to prevent comparisons between different LADs.

CTCF binds the genome in an oriented manner due to a non-palindromic motif. Loops primarily occur between two convergent CTCF motifs [34, 35]. To test whether LAD borders are enriched for a certain orientation, directionality of CTCF binding sites was assigned based on the orientation of its motif. We observe that LAD borders are enriched for CTCF binding sites with both motif orientations (Fig 1A), and therefore are likely involved in DNA looping towards (*inwards*) and away from (*outwards*) LADs.

CTCF is thus positioned at a subset of LAD borders, which raises the question whether these borders are different from LAD borders without CTCF. We classified all LAD borders into two categories based on nearby CTCF binding (positioned within 20kb outside borders, overlapping with the observed enrichment), which results in 42-62% being classified as “CTCF borders” for the different cell types (Fig 1B). In accordance with previous observations [1], LAD borders are also enriched for transcription, indicated by a local enrichment of transcription start sites (TSS) and transcription end sites (TES) for genes transcribing from and towards the LAD, respectively (Fig S1B). Active genes are present at LAD borders with and without CTCF binding (Fig 1B). To avoid confounding effects from transcription on LAD border positioning, we excluded borders near active genes from further analyses.

Average profiles of NL interactions, CTCF and cohesin binding and publicly available epigenetic data at mESC LAD borders reveal that borders with CTCF generally show a somewhat sharper transition between LAD and inter-LAD (iLAD), for example in NL interactions, ATAC-seq and H3K4me1 (Fig 1C). As expected, LAD borders with CTCF binding are locally enriched for RAD21 binding (a subunit of the cohesin complex). Finally, LAD borders with CTCF binding are locally enriched for H3K27me3 and depleted for H3K9me2. All of these enrichments are irrespective of the orientation of the CTCF motif (Fig S1C-D) and are mostly conserved in human H1 and HCT116 cells, although H3K27me3 enrichment in HCT116 cells is not limited to LAD borders (Fig S1E-F). Together, these data show that CTCF enrichment at LAD borders is conserved between cell types and that CTCF binding correlates with a different epigenetic makeup, and raise the question whether CTCF is involved in genome positioning at the NL.

### The majority of NL interactions are maintained upon depletion of CTCF

To test whether CTCF is directly involved in the demarcation of LAD borders, we used a mESC line with CTCF fused to an auxin inducible degron (AID) tag [32]. Addition of auxin (IAA) results in to rapid and near-complete protein depletion within several hours, allowing us to distinguish direct effects of CTCF depletion from secondary effects such as perturbed growth and cell differentiation [32]. We verified that IAA addition resulted in CTCF depletion (Fig S2A). Because the AID tag already resulted in a strong decrease of CTCF levels even in the absence of IAA (Fig S2A) we used the parental clone (PT) expressing only OsTir1 as an additional control in these experiments.

We then used our recently developed pA-DamID procedure to map NL interactions at multiple time points after addition of IAA. Compared to conventional DamID, pA-DamID can map NL interactions with an increased temporal resolution [33]. The pA-DamID tracks generated in the parental line are very similar to DamID tracks in wildtype mESCs except for a reduced dynamic range (Fig S2B-C), are highly reproducible between biological replicates (Fig S2D-E) and LAD borders called from pA-DamID tracks are equally enriched for CTCF binding (Fig S2F-G). Thus, the tracks generated with pA-DamID are in accordance with the DamID tracks, and can be used to generate “snapshots” of NL interactions after protein depletion. To account for differences in dynamic range present between individual replicates and conditions, we converted log_2_-ratios of LaminB1 / Dam-only to z-scores (Fig S2H). The z-scores were averaged between biological replicates for downstream analyses [33].

NL interaction patterns are largely maintained after depletion of CTCF, but there are some small differences that slowly increase over time (Fig 1D). As prolonged CTCF depletion can induce secondary effects in gene regulation and genome organization, we focused the following analyses on the early time points (6h and 24h). The average profile of NL interactions across all combined LAD borders does not reveal large differences between CTCF-marked and -unmarked LAD borders in NL interactions (Fig 1E). Similarly, we observe virtually no shift of LAD borders (which would point to expansion or contraction of LADs) after CTCF depletion (Fig 1F). However, compared to unmarked borders, we do observe a small but noticeable gain in NL interactions in the iLAD flanking CTCF-marked LAD borders (Fig 1E, red arrows), precisely coinciding with the location of the CTCF binding sites (Fig 1A). This change is already present after 6 hours, suggesting that this is a direct effect of CTCF depletion. Similar results are obtained for both CTCF orientations, despite increased variation due to further sub-grouping of the borders (Fig S2I). These data indicate that CTCF contributes to the sharpening of LAD borders, by counteracting DNA - NL contacts just outside of the borders. However, we observe no dramatic change in LAD border positions, indicating that CTCF acts in parallel with other mechanisms.

### NL detachment at CTCF LAD borders is mediated by CTCF and cohesin

The looping function of CTCF is tightly linked to cohesin, a ring-shaped protein complex involved in the formation of long-range DNA contacts. Cohesin is stalled at CTCF binding sites in the genome, and in turn is released from DNA by WAPL [reviewed in 36]. We observed that LAD borders with CTCF binding are enriched for RAD21 (Fig 1C). Moreover, we find that CTCF-marked borders coincide more frequently with loop anchors than borders without CTCF [37] (Fig 2A). These data together with the observation that WAPL knockout cells have fragmented LADs [25] suggest that cohesin may also be involved in LAD border organization. To directly test this, we then mapped NL interactions in mESCs after rapid cohesin depletion (AID-tagged RAD21) [38], cohesin stabilization (AID-tagged WAPL) [26] and a combination of CTCF depletion and cohesin stabilization (AID-tagged CTCF and WAPL) [39] (Fig S2A).

**Figure 2.**
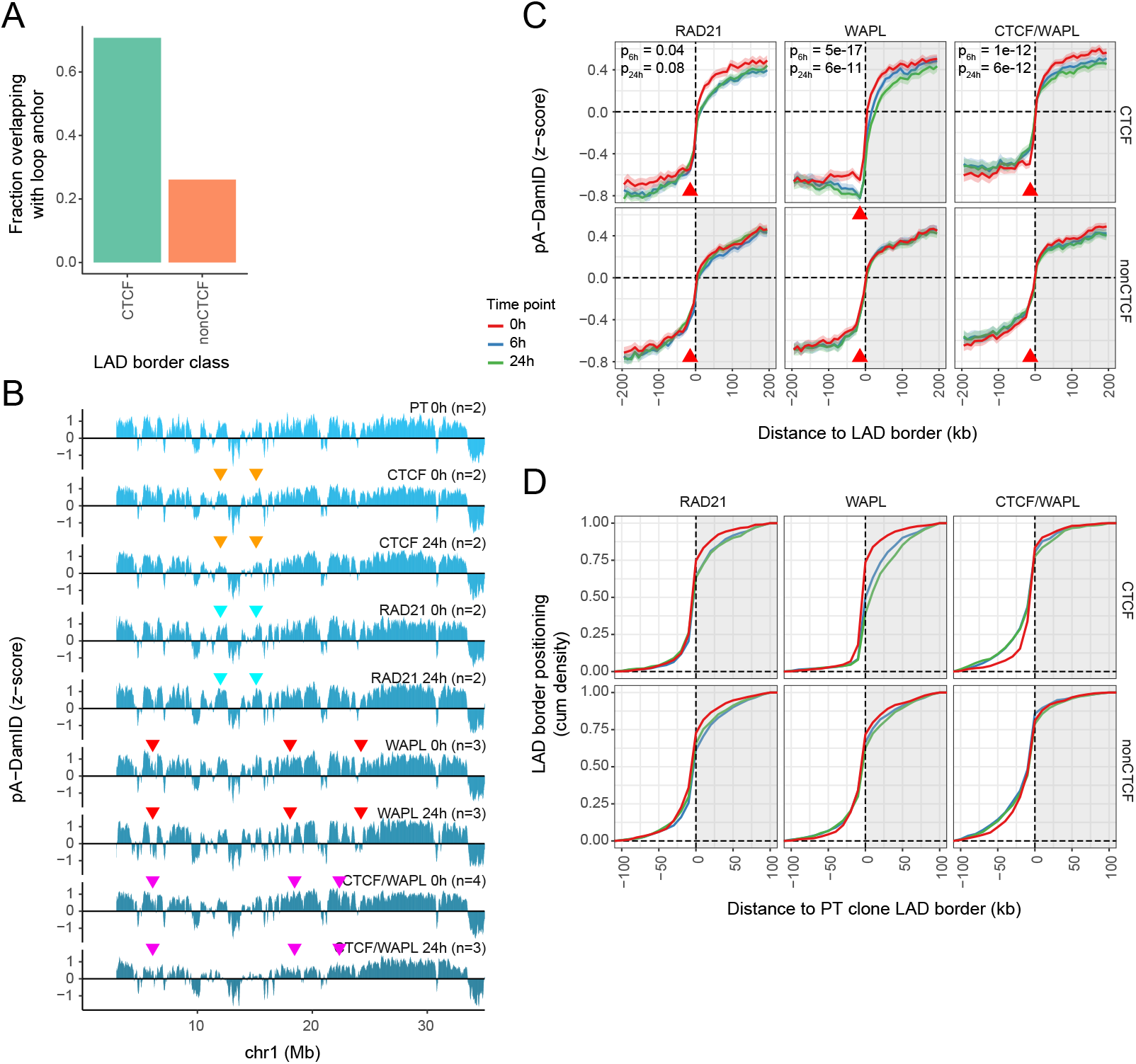
CTCF and cohesin mediate local detachment at CTCF-marked LAD borders. **(A)** Bar plot showing the fraction of LAD borders within 20 kb from a loop anchor as defined by [37]. **(B)** LaminB1 pA-DamID z-scores along a representative chromosome for all AID-tagged mESC lines without protein depletion (0h) and following 24h of IAA-induced protein depletion. Data are averages of *n* biological replicates. The colored arrows highlight example regions with deviating LaminB1 signal over time and are described in more detail in the main text. **(C)** Average LaminB1 pA-DamID z-scores around LAD borders for protein depletion time courses (0h, 6h, 24h) for mESCs with AID-tagged RAD21, WAPL and a combination of CTCF and WAPL, as described in (Fig 1E). **(D)** Cumulative density profile of LAD border positioning relative to LAD borders in the PT clone, as described in (Fig 1F).

Similar to CTCF, these depletion experiments do not affect global patterns of NL interactions, although again various small changes can be seen (Fig 2B). Intriguingly, the changes differ strongly depending on the protein depleted. For example, RAD21 depletion results in changes that are sometimes opposite of those after CTCF depletion (orange and blue arrows). WAPL depletion rapidly fragments some LADs (red arrows), similar to a prolonged loss of WAPL [25]. A double depletion of WAPL and CTCF prevents LAD fragmentation, but does result in perturbed NL interactions for some LADs (pink arrows). All these changes that can be seen by visual inspection are likely caused by a combination of various mechanisms, that we will discuss individually below.

Average profiles of NL interactions at LAD borders show that RAD21 depletion results in a small decrease in NL interactions around LAD borders marked by CTCF, both outside and inside the LAD (Fig 2C, left panels). This likely reflects a global change in NL interactions rather than a local effect of cohesin at the LAD border. We do not observe a specific shift of CTCF-marked LAD borders compared to unmarked borders (Fig 2D). A slight loss of the local dip in the average NL interaction profile at the position of CTCF binding sites (Fig 2C, red arrows) seems to occur after cohesin depletion, but this is largely masked by the overall decrease in NL interactions at CTCF-marked LAD borders.

More pronounced effects are seen after WAPL depletion (Fig 2C, middle panels). The local dip in NL interactions just outside of CTCF-marked borders becomes much more pronounced, while these borders shift on average about 10-50 kb inwards (Fig 2D). Remarkably, double depletion of WAPL and CTCF almost completely reverts this effect, underscoring that CTCF is required for cohesin-mediated local detachment (Fig 2C, right panels). Under this double depletion condition, LAD borders with CTCF binding sites tend to shift slightly towards the iLAD (Fig 2D).

Similar results are obtained for LAD borders bound by CTCF in either orientation, although the effect sizes are small for borders with outwards oriented CTCF (Fig S3A). This could be caused by an orientation effect of CTCF, but could also be due to the lower local enrichment of outwards-oriented CTCF motifs at LAD borders (Fig S2E). We note that all observed changes in pA-DamID signals discussed above are caused by a change in LaminB1 reads rather than Dam control reads, illustrating that the observed differences are not due to changes in DNA accessibility (Fig S3B). We conclude that CTCF and cohesin can promote local detachment of DNA from the NL near LAD borders, which may help to reinforce these borders.

### Cohesin-mediated local detachment of CTCF-bound chromatin from the NL inside LADs

The data presented so far indicate that chromatin bound by CTCF can locally detach DNA from the NL at LAD borders. This raises the question whether this is specific for LAD borders, or also holds true for CTCF binding sites within LADs. Small euchromatin islands in large heterochromatin domains (typically overlapping with LADs [5]) are often devoid of a TSS and enriched for active marks (DNA accessibility and H3K9ac) and binding of CTCF [40]. However, it is unclear whether CTCF is required to locally escape from the NL at these sites.

Consistent with previous work [11, 14, 41, 42], actively transcribed genes inside LADs locally detach from the NL in mESC cells (Fig 3A). To avoid confounding effects from this expression-mediated detachment, we focused on CTCF binding sites that are positioned at least 100 kb away from active genes. We find that CTCF sites inside LADs moderately detach from the NL compared to the surrounding regions (Fig 3A). Similar to LAD borders, CTCF binding in LADs is correlated with features of open chromatin (ATAC, H3K4me1 and H3K27ac), although to a much lesser extent than active genes. CTCF occupancy also coincides with a local decrease and increase of H3K9me2 and H3K27me3, respectively (Fig 3B). These data are in accordance with our observations at LAD borders.

**Figure 3.**
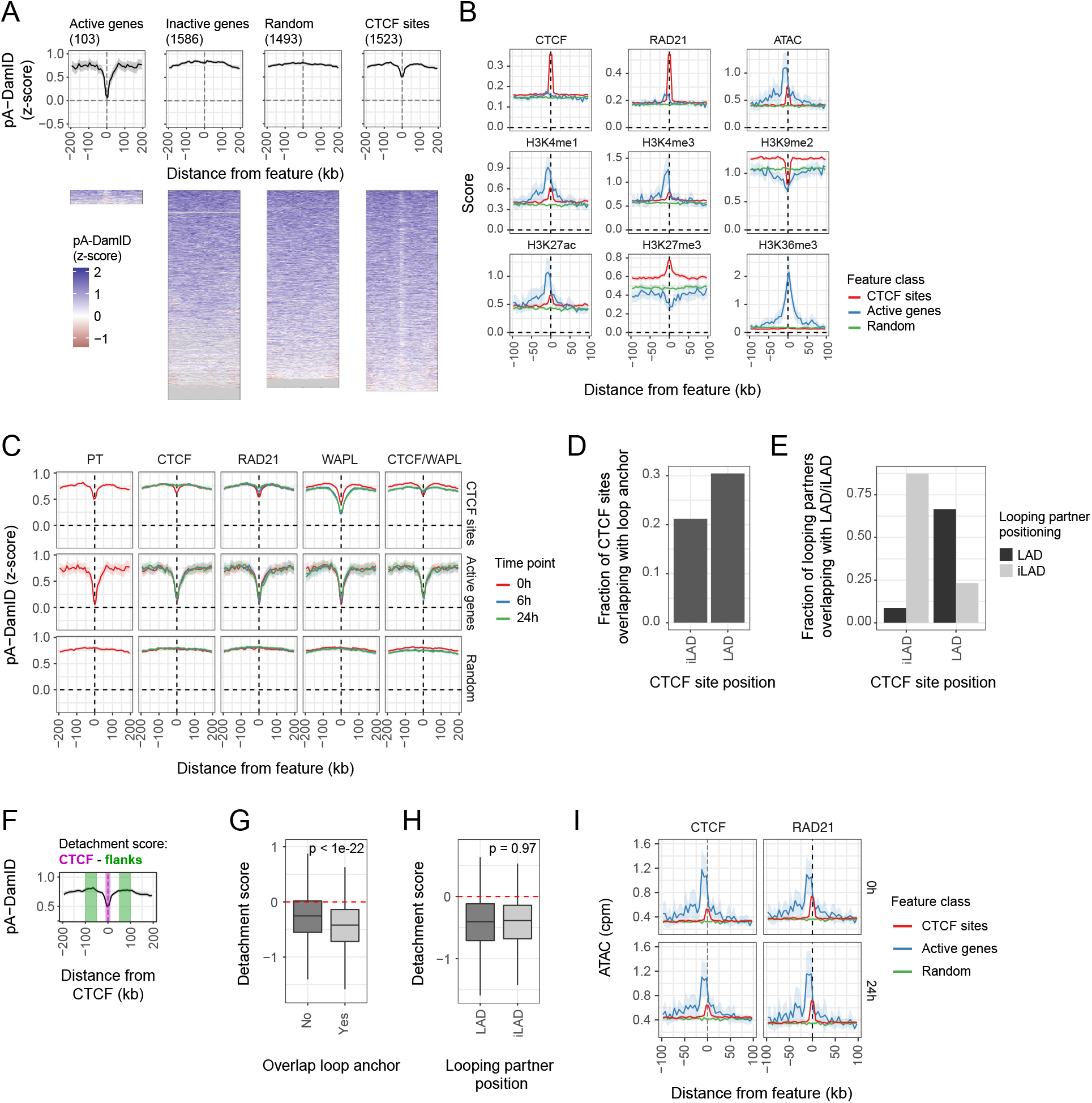
CTCF binding sites locally detach from the NL by cohesin-mediated long-range DNA interactions. **(A)** LaminB1 pA-DamID z-scores around active and inactive genes, CTCF binding sites and random locations in LADs in the mESC PT clone, visualized as average signal with 95%-confidence interval of the mean (top panel) and heatmap (bottom panel). In the heatmap, rows represent individual sites and missing data is shown as grey tiles. Genes are filtered to be positioned at least 100 kb from CTCF binding sites, and vice-versa for CTCF binding sites. **(B)** Average scores around CTCF binding sites, active genes and random locations in LADs for the epigenetic data sets, as described in (Fig 1D). **(C)** Average LaminB1 signals as in (A) following a protein depletion time course (0h, 6h and 24h) of CTCF, RAD21, WAPL and a combination of CTCF and WAPL. **(D)** Similar plots as (Fig 2A), showing the fraction of CTCF binding sites in iLADs and LADs that overlap with loop anchors. **(E)** CTCF binding sites that overlap with loop anchors are classified based on LAD overlap. The bar plot shows the positioning of their looping partners in iLAD or LAD regions. **(F)** Illustration of the detachment score, defined as the difference between the LaminB1 score at the CTCF binding site and its 100 kb-flanking region. **G)** Boxplots showing the distribution of the detachment score (F) for CTCF binding sites, separated by loop anchor overlap. CTCF binding sites within 100kb from LAD borders are excluded. A Wilcoxon test was used to test for statistical significance. **(H)** Boxplots showing the detachment score for CTCF binding sites that overlap with loop anchors (G), further separated by positioning of the looping partner. **(I)** Average ATAC-seq signals as in (B) for CTCF-AID and RAD21-AID cell lines without (0h) and with (24h) IAA-induced protein depletion.

Average NL interaction profiles across active genes, CTCF sites and random LAD locations shows that CTCF and cohesin are required for local detachment of CTCF sites within LADs (Fig 3C). WAPL depletion increases this local detachment of CTCF sites in a CTCF-dependent manner (Fig 3C). There is no change in NL interaction for active genes after any protein depletion (Fig 3C). Together, these data indicate that CTCF- and cohesin-mediated detachment from the NL is not limited to LAD borders, and that this involves a mechanism that is distinct from transcription-mediated detachment.

Next, we tried to better understand local detachment of CTCF binding sites from the NL. We hypothesized that CTCF and cohesin together mediate long-range DNA interactions of CTCF binding sites with non-LAD regions, which then compete with NL contacts. Both within and outside LADs, CTCF binding sites frequently overlap with loop anchors (Fig 3D). Loops are often formed within iLADs and within LADs, with limited cross-talk (Fig 3E). We calculated a detachment score for every CTCF binding site in LADs, defined as the difference in NL interactions between the CTCF binding site and its flanking 100kb regions (Fig 3F). In accordance with our hypothesis, local detachment is correlated with the presence of a loop anchor (Fig 3G), but surprisingly there is no difference when the looping partner is positioned outside of a LAD compared to within a LAD (Fig 3H). This suggests that long-range DNA interactions by themselves are enough to locally detach the CTCF site from the NL, and thus do not specifically require looping partners in the nuclear interior.

### Chromatin accessibility at CTCF binding sites is independent of NL detachment

Local NL detachment at CTCF binding sites with active chromatin features is thus dependent on CTCF and cohesin. This raises the question whether the detachment is required for the active chromatin features. To address this question, we generated ATAC-seq data to determine DNA accessibility [43] – one of the features enriched at CTCF binding sites – after depletion of CTCF. The results show that CTCF-mediated detachment is not required to maintain DNA accessibility (Fig 3I). We obtained similar results after depletion of RAD21 (Fig 3I). Combined, we conclude that CTCF-bound chromatin locally detaches from the NL, but that this is not required to maintain an accessible chromatin state.

### Architectural proteins control genome-wide patterns of NL interactions

Up to this point, we analyzed local changes of NL interactions at CTCF binding sites. Next, we searched for more global effects of CTCF and cohesin on NL interactions, for example involving entire LADs or large chromosomal regions. Towards this goal we calculated mean pA-DamID scores on a consensus set of LADs across all conditions, and then determined the changes in scores in each depletion experiment (see Methods, Fig S4). We observed the smallest changes in LAD scores upon CTCF depletion and the biggest changes upon the double depletion of WAPL and CTCF (Fig 4A). Consistent with visual observations of the chromosome profiles illustrated previously (Fig 2C), LAD-wide changes in NL interactions are mostly uncorrelated between the depleted proteins, except for WAPL and the double combination of CTCF and WAPL (Fig 4B).

**Figure 4.**
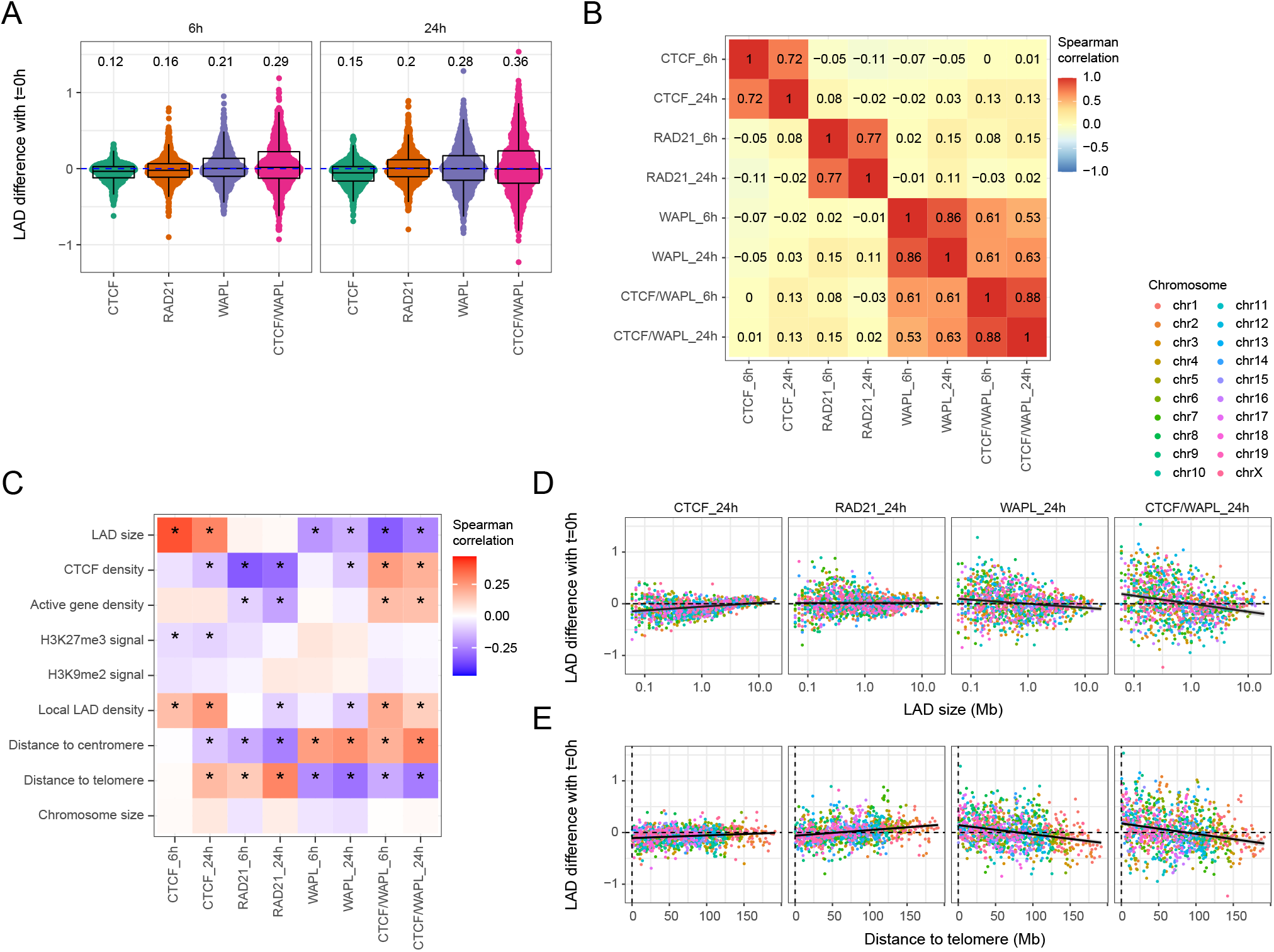
Cohesin dynamics affect chromosomal NL interaction patterns. **(A)** Quantification of LAD differences in LaminB1 pA-DamID z-scores over time (Fig S4). The number shows the coefficient of variation. **(B)** Heatmap showing Spearman correlation coefficients between the LAD differences following protein depletions. LAD differences from the same protein depletion experiment are not independent as these compare differences relative to the same starting point, and are thus inflated. **(C)** Heatmap showing Spearman correlation coefficients between LAD characteristics and differences in LAD scores. An asterisk marks a significant correlation (p < 0.05, using ‘cor.test()’ in R) after Benjamini-Hochberg multiple testing correction. The LAD characteristics are defined as follows: LAD size (log_2_ bp), CTCF binding and active gene density (count / Mb), average H3K27me3 and H3K9me2 signal, local LAD density (fraction of the surrounding 10Mb classified as LAD), distance to chromosome ends (Mb) and size of the corresponding chromosome (Mb). **(D-E)** Scatterplots of LAD differences over time versus LAD size (D) and distance to telomere (E), colored by chromosome. The black line represents a fitted linear model.

We then compared LAD-wide changes in NL interactions with several features that describe LADs and their chromosomal context: LAD size, density of CTCF binding and active genes, mean signal for the repressive histone modifications H3K27me3 and H3K9me2, local LAD density, distance to centromeres and telomeres, and chromosome size. This analysis reveals multiple significant correlations, that strongly differ depending on the depleted proteins (Fig 4C). For example, depletion of CTCF preferentially results in decreased NL interactions of small LADs, while loss of WAPL preferentially reduces NL interactions of large LADs (Fig 4D). Furthermore, the effects of RAD21 and WAPL on LAD-wide NL interactions both depend on the chromosomal position (i.e., distance to telomere and centromere), but in opposite orientation. This effect appears roughly linear with distance to the telomeres and is present across all chromosomes (Fig 4E). Chromatin nearby telomeres is enriched at the NL in early G1 cells and slowly loses interactions with the NL in interphase [33]. Possibly stabilized cohesin somehow prevents this normal maturation of NL interactions. We also found various correlations with local LAD density, CTCF density, and gene density (Fig 4C). While these intriguing global links are currently difficult to interpret mechanistically, they illustrate that the CTCF/cohesin machinery affects NL interactions not only locally, but also at the scale of entire chromosomes. Interestingly, we found only few and very modest significant correlations with H3K9me2 and H3K27me3, suggesting that the effects of perturbed cohesin dynamics are largely independent of these histone modifications.

### Genome-wide changes in NL interactions are not mediated by transcription

Altered genome positioning at the NL is strongly correlated with transcriptional changes during differentiation [6]. This is particularly relevant here, as depletions of WAPL and RAD21 induce a differentiation-like phenotype [26]. To test whether the genome-wide effects on NL interactions are mediated by transcriptional changes, we analyzed RNA-seq data after all protein depletion experiments.

Principal component analysis (PCA) (Fig S5A) and differential expression analysis (Fig 5A) indicate that all four protein depletions have a partial overlap in their effect on gene expression, although the effects of RAD21 depletion and the double CTCF/WAPL depletion are more pronounced than those of individual depletion of CTCF or WAPL. We found that genes known to be differentially expressed genes during mESC differentiation into neural precursor cells [44] are significantly affected after all depletions (Fig S5B). This implies that both CTCF and dynamic cohesin are required to maintain a normal gene regulation program and prevent mESC differentiation [26]. However, we found that LADs with up- and downregulated genes do not consistently exhibit decreased and increased NL interactions, respectively (Fig 5B). This is particularly true for LADs with early changes in gene expression (after 24h of RAD21 and CTCF+WAPL depletions). We obtain similar results with nascent transcription data following WAPL depletion (Fig S5C) [26]. We conclude that the genome-wide changes in NL interactions are generally not mediated by transcriptional changes induced by the protein depletions.

**Figure 5.**
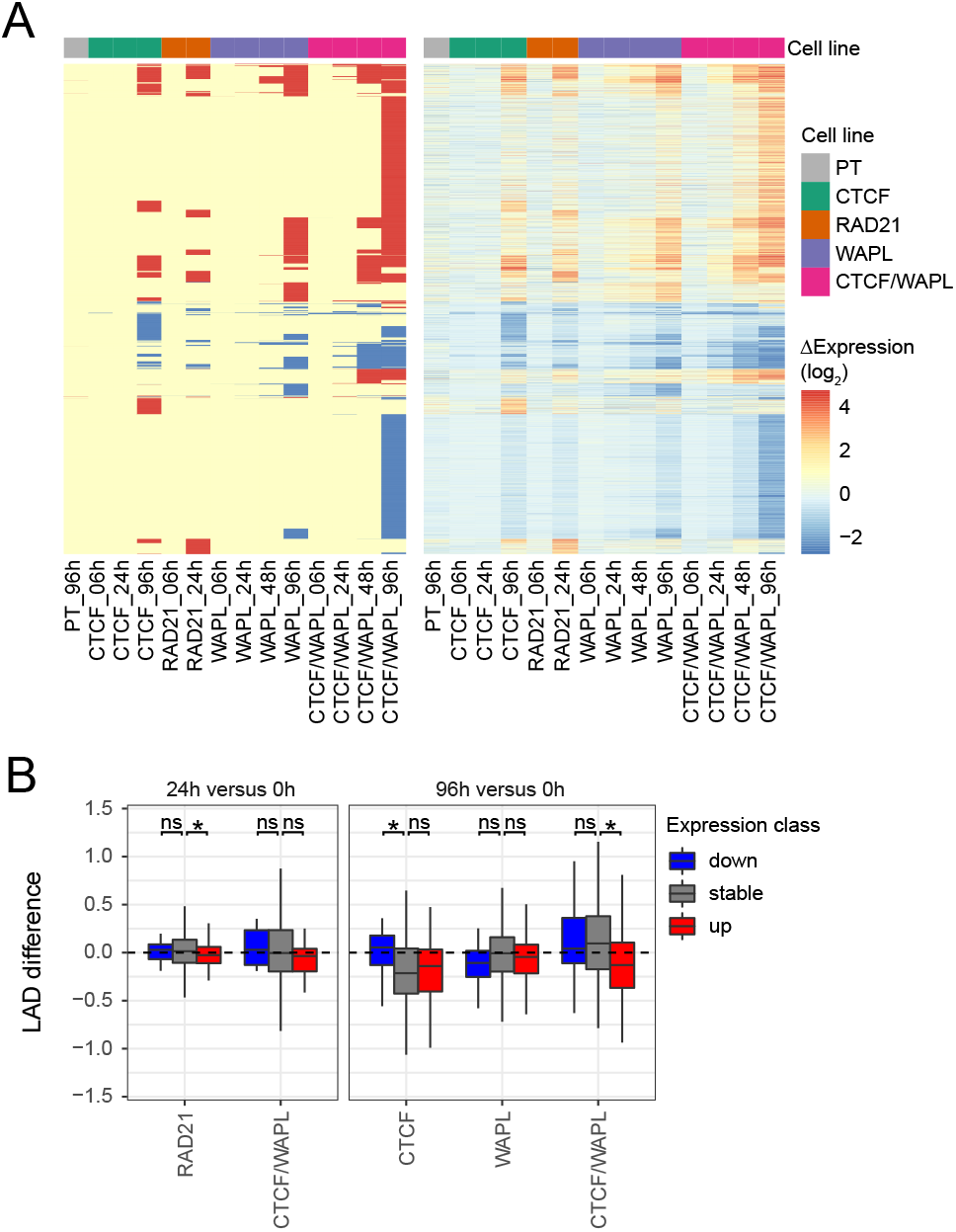
LAD dynamics are not caused by transcriptional effects of CTCF and cohesin perturbation. **(A)** Overview of RNA-seq results for time courses of CTCF depletion and cohesin perturbation, showing differentially expressed genes (left panel) and log_2_-fold changes (right panel) compared to 0h. RNA-seq data in PT, RAD21 and WAPL clones are from [26]. **(B)** Distribution of LAD differences classified by the presence of up- and down-regulated genes. LADs overlapping both up- and down-regulated genes were excluded. Only conditions with at least 50 differentially expressed genes and available gene expression and pA-DamID data are shown. A Wilcoxon test was used to test for statistical significance, followed by Benjamini-Hochberg multiple testing correction (an asterisk denotes a p-value < 0.05).

We also asked whether LAD genes are specifically affected following cohesin perturbations. Most LAD genes are not expressed and their promoters are also not expressed when placed in a neutral chromatin environment [45]. To remove such unresponsive genes we required LAD genes to have a minimum expression level in at least one of the conditions, and selected a matching set of iLAD genes with a similar distribution of expression levels as these LAD genes (Fig S5D). For all depletion experiments, differentially expressed genes are not significantly enriched in LADs (Fig S5E). We conclude that CTCF and cohesin can affect gene regulation, but not preferentially in LADs.

### NL interactions are not dependent on H3K27me3

We find that H3K27me3 is locally enriched near CTCF binding sites positioned at LAD borders and within LADs. This is of interest, because in human fibroblasts CTCF and EZH2 (methyltransferase of H3K27me3) were both reported to be involved in peripheral positioning of LAD fragments [12]. Furthermore, CTCF binding sites that overlap with H3K27me3 domains show a local increase in H3K27me3 signal after CTCF depletion in mESC cells [32]. Combined, these data may signify that a loss of CTCF is compensated by locally increased H3K27me3, and that a double depletion of CTCF and H3K27me3 is required to perturb LAD border positioning.

To test this hypothesis, we used two H3K27me3 methyltransferase inhibitors (GSK126 and EED226) that induce a near-complete loss of this mark (Fig S6A-B), and profiled NL interactions with and without CTCF depletion (Fig 6A). H3K27me3 inhibition by itself does not affect global genome positioning at the NL (Fig S6C) or specifically at LAD borders with CTCF binding (Fig 6B, top panels). Furthermore, a double depletion of H3K27me3 and CTCF has no additional effects besides those previously observed upon CTCF depletion alone (Fig 6B**, C**).

**Figure 6.**
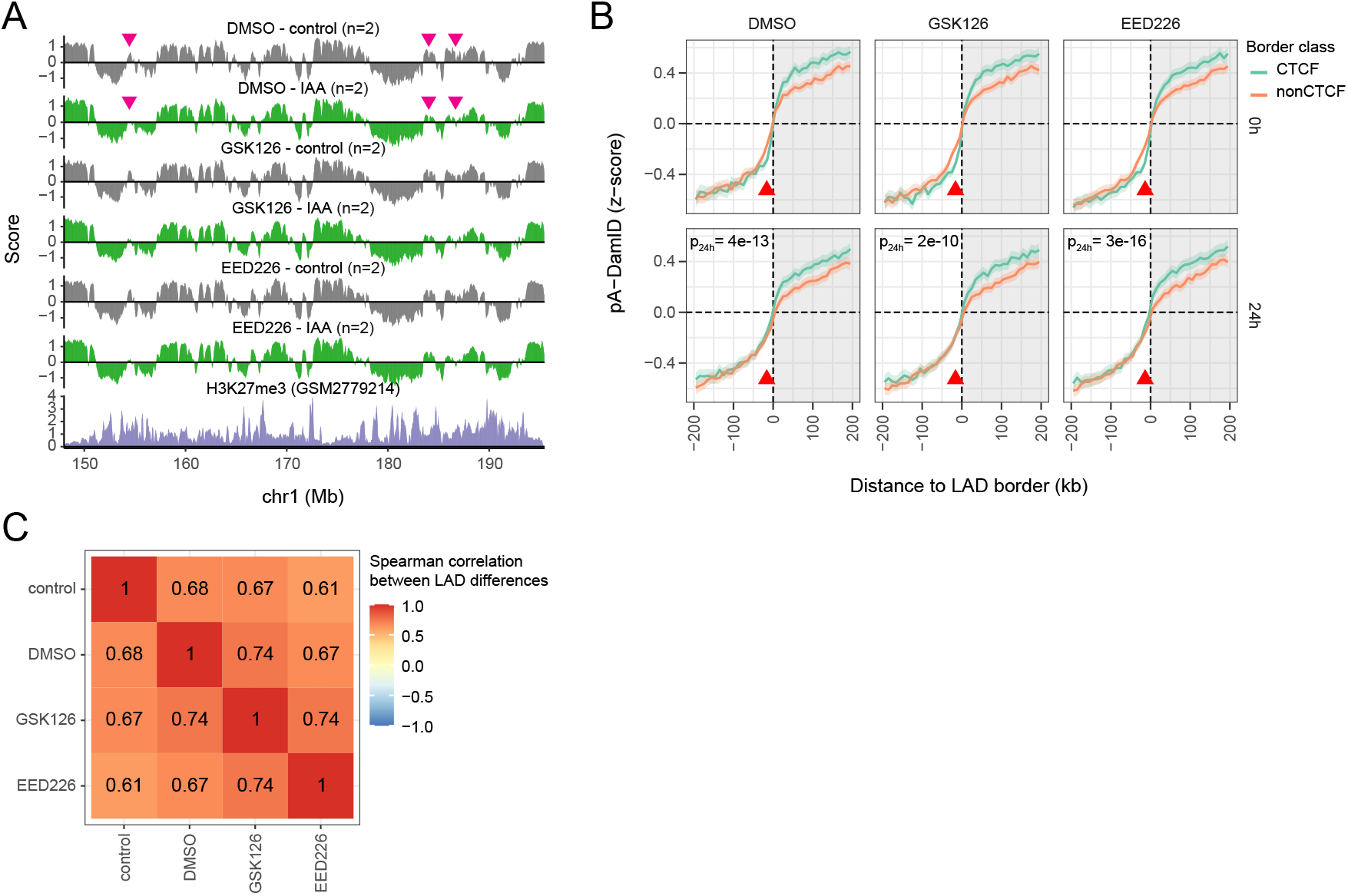
NL interactions are independent from H3K27me3 in mESCs. **(A)** Profile of LaminB1 pA-DamID z-scores along a representative genomic locus for CTCF-AID mESCs treated with DMSO and the H3K27me3 inhibitors GSK126 and EED226 (3 days), and with DMSO or IAA to induce CTCF depletion (last 24 h). Data are averages of *n* biological replicates. The pink arrows highlight example regions that lose LaminB1 signal upon CTCF depletion. H3K27me3 ChIP-seq data is added for comparison [71]. **(B)** Average LaminB1 pA-DamID z-scores around LAD borders are shown for the samples in panel (A), as described in (Fig 1E). **(C)** Heatmap showing Spearman correlation coefficients between the IAA-induced LAD differences for untreated CTCF-AID cells (control), and for cells treated with DMSO and the H3K27me3 inhibitors.

We therefore conclude that there is no interplay between CTCF and H3K27me3 in genome positioning at the NL in mESCs. This result also suggests that the genome-wide correlation between NL interactions and H3K27me3 observed after CTCF depletion is not caused by the H3K27me3 marks (Fig 4C), further supporting that the effects of perturbed cohesin dynamics on NL interactions are independent from histone modifications.

## Discussion

Combinations of chromosome conformation capture methods and rapid protein depletion experiments revealed how CTCF and cohesin organize the genome in self-associating domains [24, 25, 32, 46, 47]. However, it has remained mostly unclear how these processes affect nuclear positioning of the genome relative to the nuclear lamina. Here, we show that CTCF and cohesin reinforce LAD borders by mediating local detachment from the NL, and affect genome positioning at the NL quantitatively.

### CTCF mediates local detachment from the NL

We find CTCF and cohesin enrichment at LAD borders in all tested mouse and human cell lines, indicating that this is a conserved feature of LADs [1, 12]. Our data in mESCs indicate that these proteins are not required to maintain LAD border positioning, but rather contribute to a sharper transition in NL interactions. This result complements previous work indicating that CTCF loss does not trigger spreading of H3K27me3 heterochromatin [32]. CTCF binding may still be a boundary for other types of heterochromatin, as H3K9me2 spreading was observed following CTCF depletion for a number of genes [48], but further studies are required to validate this. Furthermore, it remains to be elucidated whether there is a functional impact of a sharper transition in NL interactions.

Small euchromatin regions inside heterochromatin blocks can be divided in promoter elements and regulatory elements based on genomic features and chromatin marks, both of which are enriched for CTCF binding sites [40]. This is in accordance with our results, which indicate that isolated CTCF sites in LADs are enriched for active marks and locally detach from the NL. We show that NL detachment at these sites requires both CTCF and cohesin. Thus, besides transcription [11, 14, 41, 42], DNA looping mediated by CTCF and cohesin can contribute to local dissociation of DNA from the NL. Upon cohesin stabilization, this CTCF-mediated detachment can fully fracture LAD domains and thus establish new LAD borders [25].

Intriguingly, CTCF and cohesin are dispensable for DNA accessibility at CTCF binding sites in LADs. This is of interest, as artificial decondensation of LADs can induce repositioning from the NL, even without transcriptional activity [7, 11, 49]. While DNA accessibility does not directly mirror chromatin decondensation, it may indicate that this repositioning also depends on CTCF and cohesin. Alternatively, these observations could be explained by a mechanism independent from transcription and CTCF-mediated DNA looping to detach from the NL.

### Cohesin dynamics affect genome-wide patterns of NL interactions

Besides local effects at LAD borders and CTCF binding sites, CTCF and cohesin affect genome-wide pattern of NL positioning quantitatively. We observe the largest effects upon depletion of WAPL and combined depletion of CTCF and WAPL. While these genome-wide changes are difficult to interpret mechanistically, they reveal intriguing correlations. For example, cohesin stabilization disrupts positioning of telomeres and nearby DNA in the nuclear interior, which is a characteristic of genome reorganization in early interphase [33, 50]. While chromatin positioning at the NL is not required for genome compartmentalization [51] and we do not find specific effects on gene expression in LADs, it remains unclear whether other phenotypes such as telomere maintenance and replication timing are affected by nuclear repositioning following perturbed cohesin dynamics [8, 52].

It was previously reported that CTCF is required for peripheral positioning of integrated LAD fragments [12]. This generally contradicts with the small effects that we observe upon CTCF depletion. We speculate that this strong effect is a consequence of the small size of the LAD fragment, which in our data correlates with decreasing NL interactions. Mixing of nearby chromatin – typically inhibited by CTCF [32, 53] – may drag small, isolated LADs into the nuclear interior.

A similar dependency for peripheral position of LAD fragments was shown for EZH2, the methyltransferase responsible for H3K27me3 deposition [12]. This differs from our results, which show that inhibition of EZH2 (with GSK126) does not affect NL interactions, alone or combined with CTCF depletion. Possibly, this discrepancy reflects cell type differences in NL affinity. H3K27me3 (and presumably also EZH2) is enriched inside LADs for some cell types, such as HCT116 cells, but not for mESCs (Fig 1C, S1F). In accordance with this reasoning, previous work illustrated that peripheral positioning of H3K27me3 is a transient phenotype during the cell cycle, development and senescence [28, 29, 54].

### What then determines LAD border positioning?

Overall, our data indicate that CTCF and cohesin are not required for LAD border positioning in mESCs, but only to strengthen the borders. It thus remains to be elucidated what demarcates LADs. While we cannot rule out that an undiscovered factor is involved, it is also possible that there are no specific characteristics of LAD borders. Rather, the factors that mediate NL interactions may be tightly controlled and result in sharp LAD borders.

Constitutive LADs and their borders are conserved between human and mouse cells [55]. While the nucleotide sequences are highly divergent, both species maintain a high AT-content in these LAD domains that presumably mediates interactions with specific NL components [55]. This may involve H3K9me2, which is evolutionary conserved, retained throughout mitosis and quickly re-establishes peripheral positioning in daughter cells [9]. Moreover, similar to constitutive LADs, mESC-specific LADs have a high AT-content [55]. This is in accordance with a model that LADs in stem cells reflect a “basal” state of chromosome organization. During differentiation, the basal organization could then be rearranged by a different NL composition that interacts with chromatin marks such as H3K27me3. In this model, CTCF, cohesin and transcription are not integral to the general pattern of LAD domains. Instead, these factors can modulate NL interactions via forces, such as the formation of loops, that counteract peripheral positioning.

## Methods

### Experimental procedures

#### Cell culture

E14Tg2a mouse embryonic stem cells (mESCs) (129/Ola isogenic background) and derived clones were cultured in serum-free DMEM/F12 (Gibco) and Neurobasal (Gibco) medium (1:1), supplemented with N-2 (Gibco), B-27 (Gibco), BSA (0.05%; Gibco), 104 U of leukemia inhibitory factor (LIF; Millipore), MEK inhibitor PD0325901 (1 μM; Selleckchem), GSK-3β inhibitor CHIR99021 (3 μM; Cayman Chemical) and 1-thioglycerol (1.5 × 10−4 M; Sigma-Aldrich) on 0.1% gelatin-coated plates. Cells were passaged every 2 days. Cells were seeded and incubated overnight before starting protein depletion experiments, at the following densities: for a 96-h time course, 35,000 and 150,000 cells were seeded in 6-well and 10-cm plates, respectively; for a 24-h time course, 0.5 million and 1.5 million cells were seeded in 6-well and 10-cm plates, respectively. During these time courses, medium was refreshed or cells were split 1:10 every 2 days.

Protein depletion was induced by treating cells with a final concentration of 500 μM IAA (I5148-10G, Sigma-Aldrich). Time series experiments were performed by inducing protein degradation at different time points and collecting all samples at the end of the time course.

F121-9-CASTx129 mESC were cultured in 2i conditions according to the 4D Nucleome guidelines (https://data.4dnucleome.org/biosources/4DNSRMG5APUM/). These cells were differentiation into neural precursor cells (mNPCs) and cultured as described [6].

Cells were tested for mycoplasma every 3 months.

#### Western blots

To verify protein depletion, AID-tagged mESCs were treated for 6 h with H_2_O or IAA and harvested. To obtain the nuclear soluble fraction, cells were resuspended in 500 μL of low salt buffer (final concentration: 10 mM HEPES, 50 mM NaCl, 1 mM EDTA, 1mM DTT, and 1X Protease Inhibitor Cocktail (11697498001, Sigma-Aldrich)) and incubated on ice for 15 min, followed by adding 500 μL of low salt/0.4% NP-40 buffer to reach a final concentration of NP-40 at 0.2%. These cells were then rotated at 4 °C for 15 min, and centrifuged at 6,000 g for 10 min at 4 °C. The obtained cell pellets were resuspended in high salt buffer (final concentration: 10 mM HEPES, 420 mM NaCl, 1 mM EDTA, 10% glycerol, and 1X Protease Inhibitor Cocktail) at a density of 5 million cells per 100 μl buffer, and then rotated overnight at 4 °C. The supernatant was collected for subsequent experiments. For the whole cell lysates, the cells were lysed in RIPA lysis buffer (150 mM NaCl, 1% NP-40, 0.5% sodium deoxycholate, 0.1% SDS and 25 mM Tris (pH 7.4)).

The nuclear extracts were used for Western blot analysis of CTCF, WAPL and ACTB, and the whole cell lysates were used for Western blot analysis of RAD21 and HSP90. Precast gradient SDS-PAGE gels (NuPAGE^TM^ 4 to 12%, Bis-Tris 1.0 mm, Mini Protein Gel, 15-well, NP0323BOX, ThermoFisher SCIENTIFIC) were used to separate the proteins, which was transferred to preactivated PVDF membranes on a Trans-Blot Turbo Transfer System (Bio-Rad). The blots were incubated with primary antibodies to the following proteins overnight at 4 °C: (1) CTCF (1:1,000; 07-729, Merck Millipore), (2) WAPL (1:1,000; 16370-1-AP, Proteintech), (3) ACTB (1:5,000; ab8227, Abcam), (4) RAD21 (1:1,000; ab154769, Abcam) and (5) HSP90 (1:2,000; 13171-1-AP). After incubation, the blots were washed three times with 0.1% Tween-20 in TBS (TBST) and then incubated with secondary antibody against rabbit IgG at room temperature for 1 h, again followed by three washes with TBST.

For H3K27me3 detection, mESCs were collected and lysed in lysis buffer (Tris pH 8.0 50 mM, EDTA 1mM, SDS 1 %) and sonicated. 25 μg of total protein extracts were run on a gradient (15-4%) Polyacrylamide gel (MINIi-PROTEAN TGX Precast Gels, Biorad). Proteins were transferred on nitro-cellulose membrane and checked with Red Ponceau staining for efficient transfer. The membranes were incubated with blocking buffer (5% milk powder in TBST) for one hour at room temperature and afterwards incubated with primary antibody (anti-H3K27me3, Diagenode # C15410195, 1:1000) in blocking buffer overnight at 4 °C. Membranes were washed with TBST and secondary antibody incubation was performed for two hours at room temperature in blocking buffer.

Protein detection was performed using Biorad Clarity Max ECL substrates and a ChemiDoc MP Imaging System (Bio-Rad).

#### LaminB1 DamID and pA-DamID

LaminB1 DamID was performed as described previously [45, 56]. Briefly, F121-9-CASTx129 mESCs and mNPCs were lentiviral transduced in a 6-well plate with Dam control or (mouse) Dam-LaminB1 constructs. Dam was under control of the human PGK1 promoter and fused to a destabilization domain. Genomic DNA was isolated three days after transduction (ISOLATE II Genomic DNA kit, Bioline BIO-52067), without stabilization of the Dam fusions to keep expression at low levels. ^m6^A-marked DNA was enriched using a sequence of DpnI digestion, PCR adapter ligation, DpnII digestion and PCR amplification. Resulting amplified material was processed for high-throughput sequencing using an in-house library preparation procedure, and sequenced for single-end 140 bp reads on an Illumina HiSeq 2500. Approximately 10 million reads were sequenced for every condition.

pA-DamID LaminB1 maps were generated as described [33]. One million E14Tg2a mESCs were collected by centrifugation (500 g, 3 minutes) and washed sequentially in ice-cold PBS and digitonin wash buffer (DigWash) (20 mM HEPES-KOH pH 7.5, 150 mM NaCl, 0.5 mM spermidine, 0.02% digitonin, cOmplete Protease Inhibitor Cocktail). Cells were rotated for 2 hours at 4°C in 200 μL DigWash with 1:400 Lamin B1 antibody (Abcam, ab16048, rabbit), followed by a wash step with DigWash. This was repeated with a 1:200 pA-Dam solution (∼60 NEB Dam units), followed by 2 wash steps. Dam activity was induced by an incubation for 30 minutes at 37°C in 100 μL DigWash supplemented with 80 μM SAM while gently shaking (500 rpm). Genomic DNA was isolated and DNA was processed similar to DamID, except that the DpnII digestion was omitted and 65 bp reads were sequenced. For every condition, another 1 million cells were processed in only DigWash and during Dam activation incubated with 4 units of Dam enzyme (NEB, M0222L). This Dam control sample serves to account for DNA accessibility and amplification biases.

#### ATAC-seq

ATAC-seq libraries were prepared as previously described [57]. Cells were permeabilized and tagmented with in-house-generated Tn5 transposase, after which DNA fragments were amplified with two sequential nine-cycle PCR runs. Fragments smaller than 700 bp were selected with SPRI beads. ATAC-seq libraries were sequenced with paired-end mode using a 75-cycle kit on an Illumina NextSeq 550.

#### ChIP-seq

ChIP-seq was performed as described but with small modifications [57]. mNPCs were mixed with 10% HEK293T cells (as internal reference) and crosslinked for 10 min with 1% FA. The reaction was quenched with 2.0 M glycine. After cell lysis, a Bioruptor Plus sonication device (Diagenode) was used to fragment chromatin to approximately 300 bp fragments. CTCF antibody (07-729, Merck Millipore; 5 μL per ChIP) was coupled with Protein G beads (Thermo Fisher Scientific) and incubated with fragmented chromatin overnight at 4 °C. After washing, chromatin was eluted from the beads and crosslinking was reversed. Released DNA fragments were purified with the MinElute PCR Purification kit (Qiagen). The purified DNA fragments were prepared with the KAPA HTP Library Preparation kit (Roche) and single-end sequenced with a 65-cycle kit on an Illumina HiSeq 2500.

#### RNA-seq

RNA was isolated using a standard TRIzol RNA isolation protocol (Ambion). Cells were lysed with 1 mL of TRIzol reagent, after which 200 μL chloroform was added and the mixture was vortexed. Following centrifugation at 12,000g at for 15 min at 4 °C, the upper phase was homogenized with 0.5 mL of 100% isopropanol. Samples were incubated at room temperature for 10 min and centrifuged at 4 °C for another 10 min. The resulting RNA pellet was washed with ice-cold 75% ethanol, dried at room temperature and resuspended in RNase-free water. The isolated RNA was treated with DNase using the RNeasy Mini kit (Qiagen). RNA-seq libraries were prepared using a TruSeq Stranded RNA LT Kit (Illumina). The libraries were sequenced for single-end 65 bps reads on an Illumina HiSeq 2500.

### Computational analyses

#### DamID and pA-DamID

DamID and pA-DamID data was processed as described [45]. Briefly, the adapter sequence was trimmed with cutadapt 1.11 before mapping the remaining genomic DNA sequence to mm10 with bwa mem 0.7.17. The following steps were performed with custom R scripts. Reads with a mapping quality of at least 10 and overlapping the ends of a GATC fragment were counted in 10kb genomic bins. Counts were normalized to 1M reads per sample and a log2-ratio over the Dam control sample was calculated with a pseudocount of 1. At least 2 biological replicates were generated for every experimental condition and the average score was used for downstream analyses. Log2-ratios were converted to z-scores to correct for differences in dynamic range between experiments (Fig S2H).

#### LAD definition

LADs were determined using hidden Markov modeling on the average NL interaction profile between biological replicates (https://github.com/ gui11aume/HMMt). LADs within 50kb of each other were merged and LADs were filtered to be at least 50kb in size. For Figures 1A-C, the LAD definition based on wildtype mESC (F121-9-CASTx129 strain) LaminB1 DamID was used. For the remaining analyses the LAD definition was based on PT mESC (E14Tg2a strain) LaminB1 pA-DamID, except when otherwise stated. In order to capture newly formed LADs, a consensus LAD model between all experimental conditions was used for Figures 4 and 5 (a union set of LADs called in PT and AID-tagged cell lines up to 24h of IAA addition). A similar consensus LAD model was used for Figure 6 using CTCF-AID cells treated with DMSO and H3K27me3 inhibitors and up to 24h of IAA addition.

#### LAD border classification

LAD borders were classified as CTCF borders if a CTCF binding site was within 20 kb outside the LAD (overlapping with the enriched CTCF density). Borders within 10 kb of an active gene (FPKM > 1) were flagged and not used in downstream analyses. LAD borders were assigned a loop anchor if this was positioned within 20 kb. To determine CTCF orientation at LAD borders, the CTCF motif (JASPAR MA0139.1 [58]) was used to infer the orientation of CTCF binding sites with FIMO (MEME suite) [59]. LAD borders with a single CTCF orientation were assigned this orientation, while LAD borders near multiple CTCF orientations or unassigned CTCF binding sites were assigned “ambiguous”. Deeptools 3.5.0 [60] and custom R scripts were used to calculate and visualize scores relative to LAD borders and LAD features (i.e. active genes, CTCF binding sites).

#### Nuclear lamina detachment score

The detachment score was defined as the difference in mean NL interaction scores between the flanking region (distances between 50-100 kb, both sides) and the CTCF binding site (up to 10 kb). To prevent confounding factors for transcription and border positioning, CTCF binding sites were filtered to be at least 100 kb from active genes and LAD borders.

#### RNA-seq and TT-seq

RNA-seq reads were mapped against the mm10 reference genome with TopHat2 2.1.1 [61] and filtered for a mapping quality of at least 10. Read counts for Ensembl genes (GRCm38.92; exons only) were determined with HTSeq 0.9.1. DESeq2 1.30.1 [62] was used to call differentially expressed genes by testing for a log2-fold-difference of 0.5 with a false discovery rate of 0.05. Principal component analysis was performed with ‘plotPCA()’ from DESeq2, using the top5000 most variable genes. FPKM values were calculated with ‘fpkm()’ from DESeq2, using the combined exon length as gene length. Similar downstream processing was applied to call differential expression in the TT-seq data [26].

#### ATAC-seq

ATAC-seq reads were mapped to mm10 with bwa mem 0.7.15-r1140. SAMtools 1.9 [63] was used to filter reads for a mapping quality of at least 15 and discard optimal PCR duplicates. Genomic coverage was determined with deeptools 3.0 [60].

#### ChIP-seq

ChIP-seq data were analyzed as previously reported [26]. Reads were mapped to a concatenated reference genome (mm10 and hg19) using bowtie2 2.3.4.1 [64]. Reads were filtered for a mapping quality of at least 15 and optical PCR duplicates were removed with SAMtools 1.9. The normalization factor to scale the spike-in HEK293T reads (mapping to hg19) to 1 million reads was used as the scaling factor in deeptools 3.0 to determine genomic coverage. Peak calling was performed with MACS2 2.1.1.20160309 [65] at a q-value cutoff of 0.01.

#### External data

The external datasets that have been used in this study are listed in the supplementary information.

## Declarations

### Ethics approval and consent to participate

Not applicable.

### Consent for publication

Not applicable.

### Availability of data and materials

The genomic datasets and computational code produced in this study are available from the following repositories:

- mESC DamID data, raw and processed files: 4D Nucleome data repository (https://data.4dnucleome.org/).
- Other genomics data: GSE181693 (pA-DamID), GSE181849 (RNA-seq), GSE181846 (ATAC-seq) and GSE183958 (mNPC data).
- Computational code (will be made available before publication) : https://github.com/vansteensellab/xxx.

### Competing interests

TvS and BvS are in discussion with a company to make the pA-DamID reagents commercially available.

### Funding

Supported by NIH Common Fund “4D Nucleome” Program grant U54DK107965 (BvS), European Research Council Advanced Grant 694466 (‘GoCADiSC’) (BvS), Marie Curie Fellowship 838555 (SGM), ERC Starting Grant 637587 (‘HAP-PHEN’) (EdW), ERC Consolidator Grant 865459 (‘FuncDis3D’) (EdW), Vidi grant from the Netherlands Scientific Organization (NWO, 016.16.316) (EdW) and Veni grant from the Netherlands Scientific Organization (NWO, 016.Veni.181.014) (NQL). The Oncode Institute is partly supported by KWF Dutch Cancer Society.

### Authors’ contributions

TvS: Conceived and designed study, performed experiments, conducted majority of data analysis, wrote manuscript. NQL: Conceived and designed study, performed experiments and data analysis. SGM: Performed experiments. EdW: Designed study, supervised project. BvS: Designed study, wrote manuscript, supervised project.

## Acknowledgements

We thank the NKI Digital Microscopy, Flow Cytometry, Genomics, Protein Production, and RHPC core facilities for technical assistance, and members of our laboratories for helpful comments.

---

**Supplementary figure 1.**
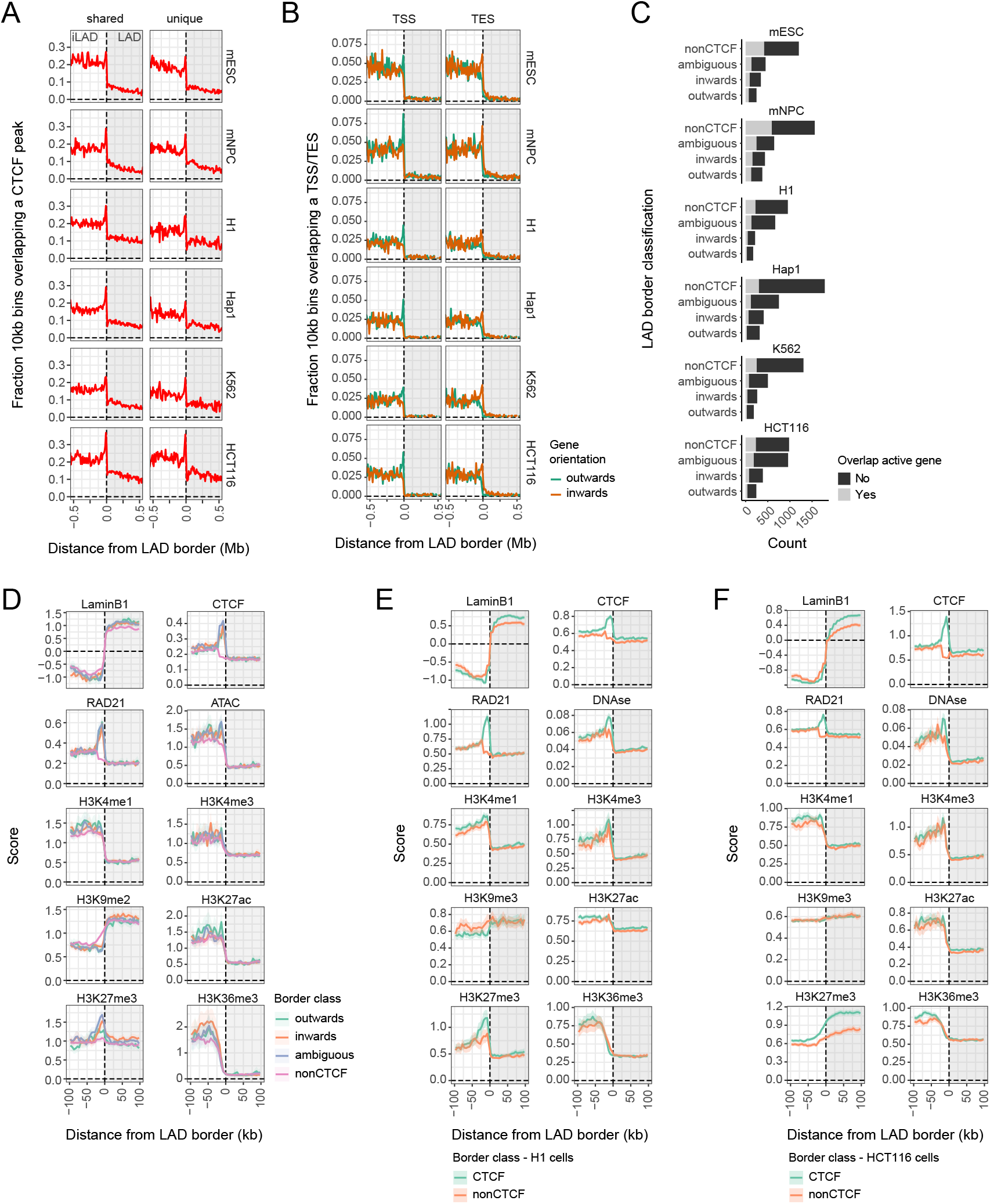
LAD border enrichment of epigenetic marks is independent of CTCF orientation and is conserved between cell types. **(A)** Profiles similar to (Fig 1A), showing CTCF enrichment around LAD borders classified on conservation between cell types. LAD borders are classified as shared between cell types when a LAD border in another cell type of the same species is within 100kb. **(B)** TSS and TES enrichment of active genes (FPKM > 1) is shown around LAD borders. Gene orientation was used to determine the orientation relative to the LAD, where inwards and outwards indicate a gene transcribing towards and away from the LAD, respectively. **(C)** Classification of LAD borders similar as (Fig 1B), but including CTCF motif orientation in the LAD border classification. LAD borders overlapping with CTCF binding sites for which the orientation could not be determined and borders overlapping with multiple orientations were classified as ambiguous LAD borders. Significantly more CTCF binding sites have been called in H1 and HCT116 cell lines (55-60×10^3^ compared to 30-45×10^3^), resulting in a larger fraction of ambiguous LAD borders for these cell lines. **(D)** Profiles similar as (Fig 1C), but using LAD borders classified with CTCF motif orientation. **(E-F)** Profiles similar as (Fig 1C), but for LAD borders in H1 (E) and HCT116 (F) cells [67]. Due to data availability and data quality, ATAC-seq was replaced by DNAse as a measure for DNA accessibility and H3K9me2 was replaced by H3K9me3 in both cell types.

**Supplementary figure 2.**
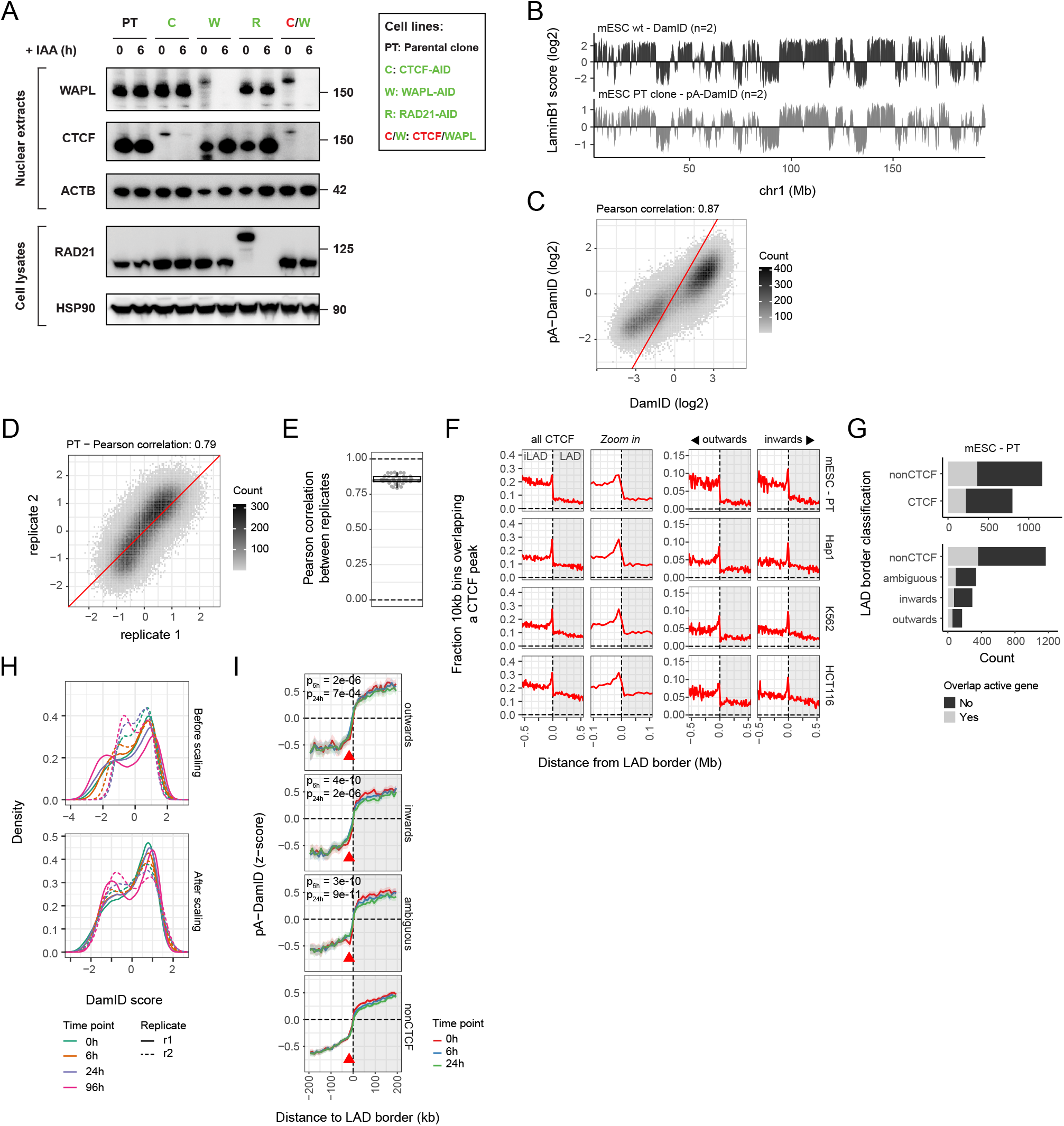
LaminB1 pA-DamID tracks recapitulate DamID data and can be used to profile NL interactions after rapid protein depletion. **(A)** Western blot analysis of WAPL, CTCF and RAD21 levels in the parental line (PT) and AID-tagged mESC clones before and after 6 hours of IAA treatment (cropped, n = 1). Green and red text indicate GFP and mCherry fusions, respectively. **(B)** Profile of LaminB1 interactions scores along a representative chromosome generated with DamID (in wildtype mESCs, F121-9 strain) and pA-DamID (in mESC PT clone, E14Tg2a strain). The log_2_-ratios of LaminB1 reads over Dam-control reads are shown for 10 kb genomic bins. Data tracks are averages of *n* biological replicates. **(C)** Scatterplot of LaminB1 interaction scores from (B) for all chromosomes. The red line represents the diagonal. **(D)** Scatterplot of LaminB1 pA-DamID interaction scores (log_2_-ratios) for two PT clone replicates. The red line represents the diagonal. **(E)** Distribution of Pearson correlation values between all biological replicate experiments described in this manuscript. **(F)** Profiles similar as (Fig 1A), showing CTCF enrichment around LAD borders based on pA-DamID data. **(G)** LAD border classification plots similar as (Fig 1B) and (Fig S1C), for LADs defined with pA-DamID in the mESC PT clone. **(H)** Data distribution of LaminB1 pA-DamID log_2_-ratios before and after conversion to z-scores. The distribution shapes are not affected by this transformation. Every line represents a single replicate. **(I)** Profiles similar as (Fig 1E), showing pA-DamID scores for LAD borders classified with CTCF motif orientation.

**Supplementary figure 3.**
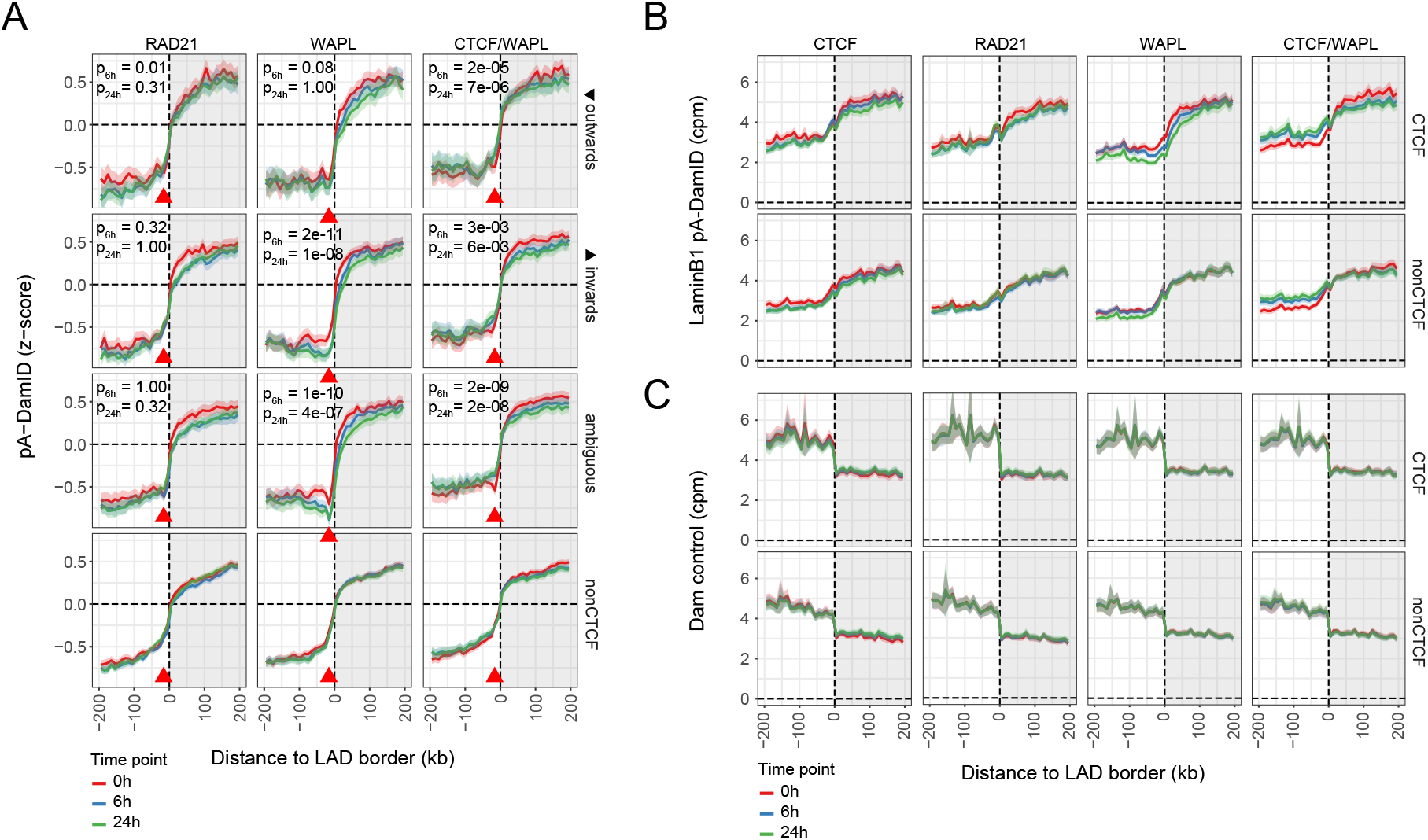
Perturbed NL interactions are limited for outwards oriented LAD borders and caused by changes in LaminB1 reads. **(A)** Profiles similar as (Fig 2D), showing pA-DamID scores for LAD borders classified with CTCF motif orientation. **(B-C)** Counts-per-million normalized (CPM) LaminB1 (B) and Dam-control (C) signals are shown around LAD borders.

**Supplementary figure 4.**
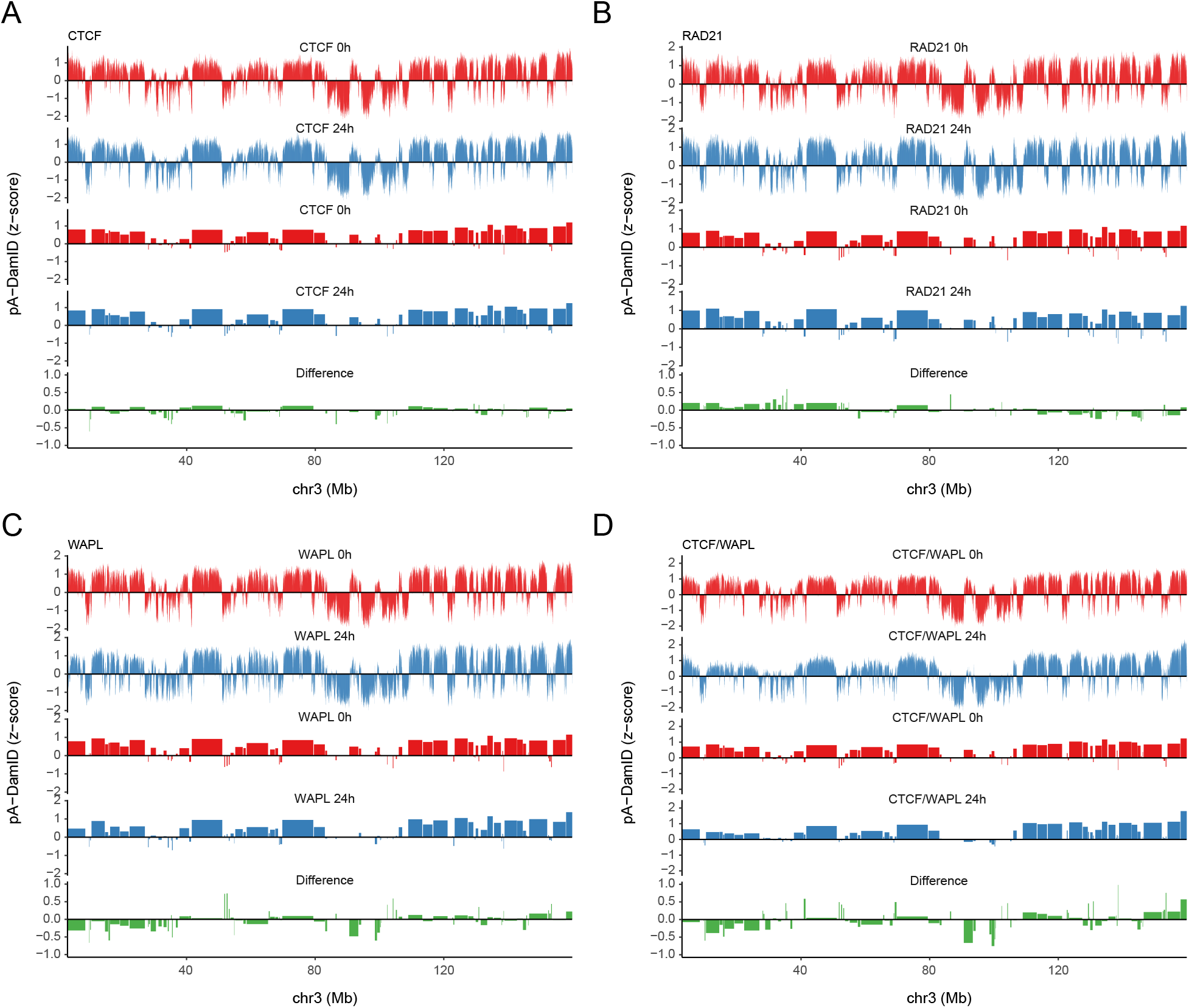
Quantification of LAD differences after CTCF depletion and cohesin perturbation. **(A-D)** Overview of the quantification of NL interactions changes along a representative chromosome, for CTCF (A), RAD21 (B), WAPL (C) and CTCF and WAPL (D) depletion experiments (tracks 1-2). A union set of LADs was created across all cell lines and conditions. The LAD score is defined as the average signal across all overlapping 10 kb bins (tracks 3-4). The difference in LAD score (track 5) was used to correlate with LAD features, such as LAD size, gene density and chromosomal positioning.

**Supplementary figure 5.**
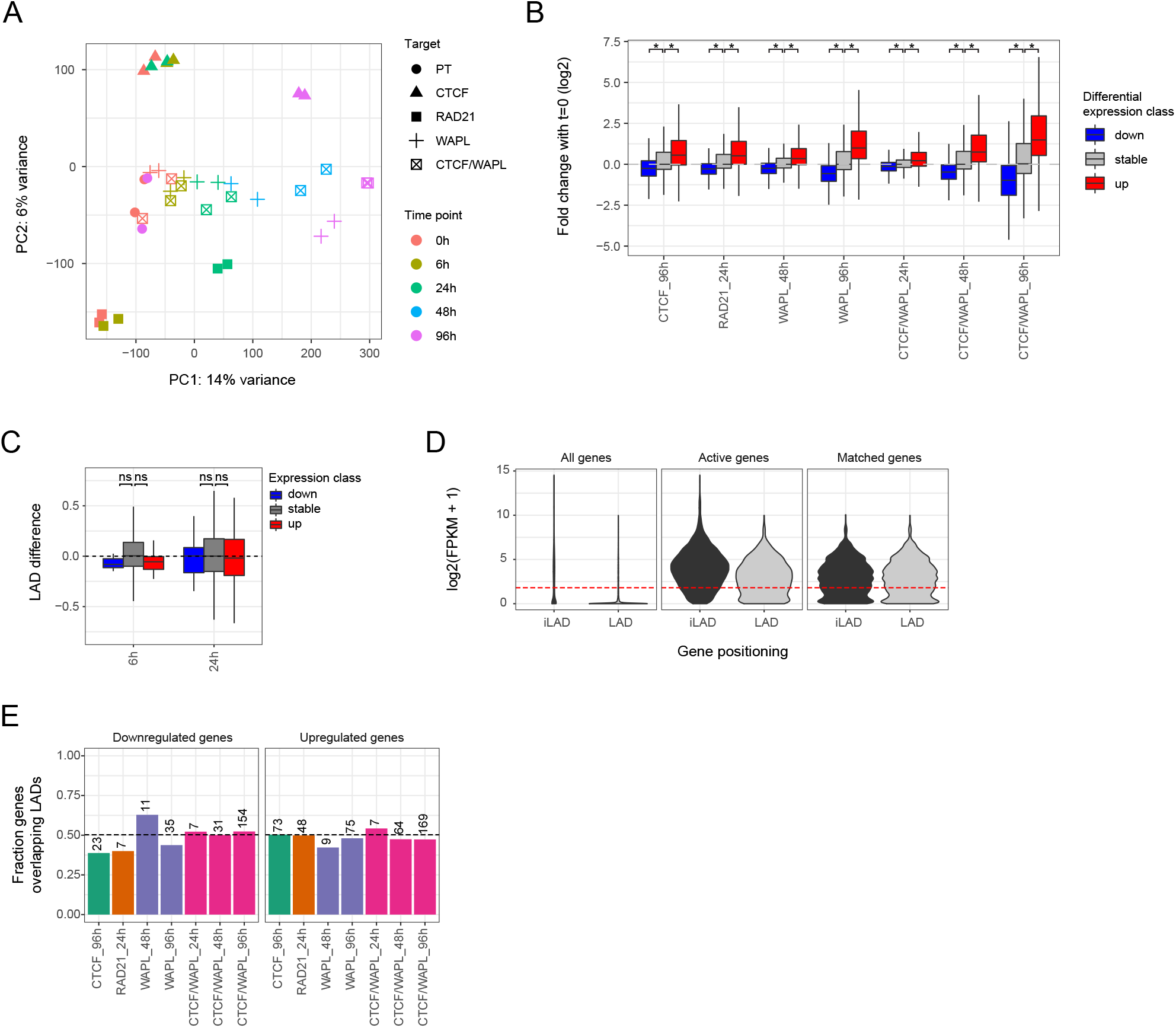
CTCF depletion and cohesin perturbation affect an overlapping gene set enriched for differentiation genes. **(A)** Principal component analysis (PCA) for all RNA-seq samples based on the top-5000 most variable genes. **(B)** Box plots showing expression fold changes (log_2_) relative to 0h of protein depletion for differentially expressed genes during mESC to mNPC differentiation [44]. Only time points with at least 50 differentially expressed genes are shown. Significance was tested for using a Wilcoxin test followed by Benjamini-Hochberg multiple testing correction. **(C)** Similar plot as (Fig 5B), but showing LAD differences classified by nascent transcription data after WAPL depletion in WAPL-AID cells. **(D)** FPKM distributions for genes positioned in iLADs and LADs. Genes were filtered to be active in at least one condition (active genes) and a random matching set of iLAD genes was selected with similar expression as the LAD genes (matched genes). 10 random sets were taken for a robust comparison. **(E)** Fraction of differentially expressed genes positioned in LADs for using an equal-sized matched expression set of LAD and iLAD genes (D). A fraction of 0.5 indicates that LAD and iLAD genes are equally affected by protein depletions. The number denotes the number of differentially expressed LAD genes.

**Supplementary figure 6.**
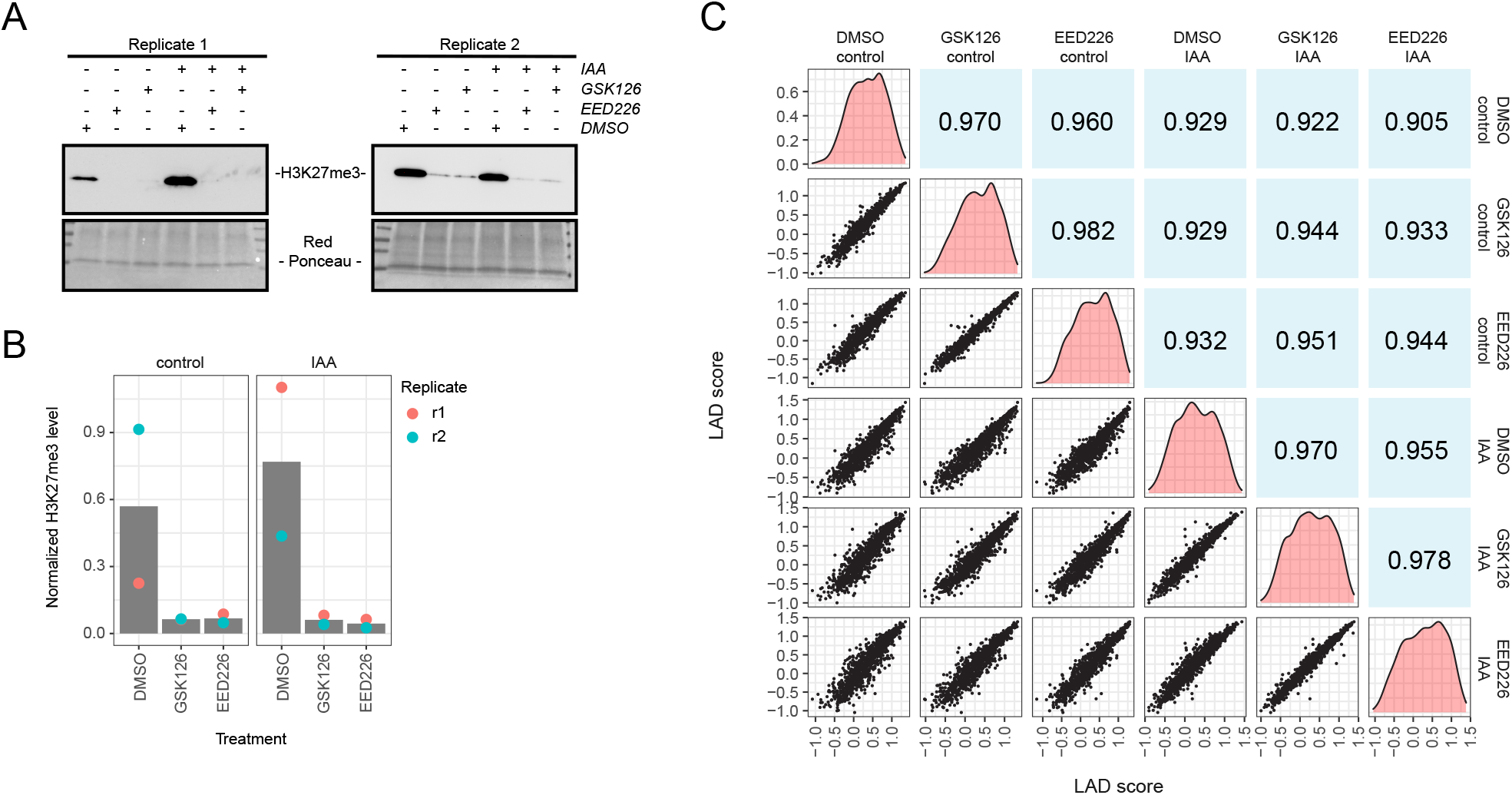
H3K27me3 depletion does not affect genome-wide NL interactions. **(A)** Western blot analysis of H3K27me3 levels in CTCF-AID mESCs treated for 3 days with the H3K27me3 inhibitors GSK126 (1 µM final concentration from 1 mM stock solution in DMSO) and EED226 (10 µM final concentration from 20 mM stock solution in DMSO) or a similar amount of DMSO as control, with and without 24 hours of IAA addition (cropped, n = 2). **(B)** Quantification of H3K27me3 levels compared to loaded protein (Ponceau staining). **(C)** Correlation matrix of LAD scores for the conditions listed in (A), showing scatter plots (bottom-left panels), data distributions (diagonal panels) and Pearson correlations (top-right panels) for all comparisons. Note that the variation upon IAA addition is larger than the variation induced by H3K27me3 depletion.

