## Supplementary Information for "CTCF and cohesin promote focal detachment of DNA from the nuclear lamina"

**Data sets used**

| Data type | Cell type | Target | Reference | ID |
|---|---|---|---|---|
| DamID data to define LADs | | | | |
| DamID | mESC | LaminB1 | 4D Nucleome; this work | 4DNESYPMROEJ |
| DamID | mNPC | LaminB1 | This work | GSE183958 |
| DamID | H1 | LaminB1 | 4D Nucleome; this work | 4DNESXKBPZKQ |
| DamID | Hap1 | LaminB1 | 4D Nucleome; van Schaik, 2020 | 4DNESUK5H9Y8 |
| DamID | K562 | LaminB1 | 4D Nucleome; van Schaik, 2020 | 4DNESTAJJM3X |
| DamID | HCT116 | LaminB1 | 4D Nucleome; van Schaik, 2020 | 4DNES24XA7U8 |
| CTCF ChIP-seq to determine LAD border enrichment | | | | |
| ChIP | mESC | CTCF | Liu, 2021 | GSM3992899 |
| ChIP | mNPC | CTCF | This work | GSE183958 |
| ChIP | H1 | CTCF | ENCODE | ENCSR000BNH |
| ChIP | Hap1 | CTCF | Haarhuis, 2017 | GSM2493878 |
| ChIPmentation | K562 | CTCF | Schmidl, 2015 | SRR2085872 |
| ChIP | HCT116 | CTCF | ENCODE | ENCSR240PRQ |
| pA-DamID data generate after acute protein depletion | | | | |
| pA-DamID | mESC + various conditions | LaminB1 | This work | GSE181693 |
| pA-DamID | Hap1 | LaminB1 | 4D Nucleome; van Schaik, 2020 | 4DNESFWILAC9 |
| pA-DamID | K562 | LaminB1 | 4D Nucleome; van Schaik, 2020 | 4DNESUMP6SS1 |
| pA-DamID | HCT116 | LaminB1 | 4D Nucleome; van Schaik, 2020 | 4DNESWB729QB |
| RNA-seq generated after acute protein depletion | | | | |
| RNA-seq | mESC PT | | Liu, 2021 | GSE135180 |
| RNA-seq | CTCF-AID | | This work | GSE181849 |
| RNA-seq | RAD21-AID | | Liu, 2021 | GSE135180 |
| RNA-seq | WAPL-AID | | Liu, 2021 | GSE135180 |
| RNA-seq | CTCF/WAPL-AID | | This work | GSE181849 |
| ATAC-seq generated after acute protein depletion | | | | |
| ATAC-seq | CTCF-AID | | This work | GSE181846 |
| ATAC-seq | RAD21-AID | | This work | GSE181846 |
| mESC loop pairs | | | | |
| Hi-C | Loop pairs | | Bonev, 2017 | GSE96107 |
| mESC epigenetic data set | | | | |
| ChIP | mESC | RAD21 | Liu, 2021 | GSM3992901 |
| ATAC | mESC | | Tastemel, 2017 | GSM2651155 |
| ChIP | mESC | H3K4me1 | Joshi, 2015 | GSM1856424 |
| ChIP | mESC | H3K4me3 | Marks, 2012 | GSM590112 |
| ChIP | mESC | H3K9me2 | von Meyenn, 2016 | GSM2051618 |
| ChIP | mESC | H3K27ac | Joshi, 2015 | GSM1856426 |
| ChIP | mESC | H3K27me3 | Højfeldt, 2018 | GSM2779214 |
| ChIP | mESC | H3K36me3 | Marks, 2012 | GSM590120 |
| H1 epigenetic data set | | | | |
| ChIP | H1 | RAD21 | ENCODE | ENCSR000BLD |
| DNAse | H1 | | ENCODE | ENCSR794OFW |
| ChIP | H1 | H3K4me1 | ENCODE | ENCSR631RJR |
| ChIP | H1 | H3K4me3 | ENCODE | ENCSR019SQX |
| ChIP | H1 | H3K9me3 | ENCODE | ENCSR883AQJ |
| ChIP | H1 | H3K27ac | ENCODE | ENCSR000ANP |
| ChIP | H1 | H3K27me3 | ENCODE | ENCSR216OGD |
| ChIP | H1 | H3K36me3 | ENCODE | ENCSR476KTK |
| HCT116 epigenetic data set | | | | |
| ChIP | HCT116 | RAD21 | ENCODE | ENCSR000BSB |
| DNAse | HCT116 | | ENCODE | ENCSR000ENM |
| ChIP | HCT116 | H3K4me1 | ENCODE | ENCSR161MXP |
| ChIP | HCT116 | H3K4me3 | ENCODE | ENCSR333OPW |
| ChIP | HCT116 | H3K9me3 | ENCODE | ENCSR179BUC |
| ChIP | HCT116 | H3K27ac | ENCODE | ENCSR000EUT |
| ChIP | HCT116 | H3K27me3 | ENCODE | ENCSR810BDB |
| ChIP | HCT116 | H3K36me3 | ENCODE | ENCSR091QXP |
| RNA-seq in other cell types for active gene definition (FPKM > 1) | | | | |
| RNA-seq | mNPC | | Bonev, 2017 | GSE96107 |
| RNA-seq | H1 | | ENCODE | ENCFF000FET, ENCFF000FEU, ENCFF000FEU, ENCFF000FET, ENCFF000DJM, ENCFF000DJN, ENCFF000DJM, ENCFF000DJN, ENCFF565ZQD, ENCFF565ZQD, ENCFF953ZDW, ENCFF953ZDW, ENCFF589VNC, ENCFF608OLY, ENCFF589VNC, ENCFF247WDK, ENCFF350HDB, ENCFF199CUO, ENCFF608OLY, ENCFF687WLZ, ENCFF199CUO, ENCFF350HDB, ENCFF247WDK, ENCFF567PCA, ENCFF567PCA, ENCFF687WLZ |
| RNA-seq | K562 | | ENCODE | ENCFF001RED, ENCFF001REG, ENCFF001RWD, ENCFF001RVV, ENCFF001RWE, ENCFF001RWF, ENCFF001RDD, ENCFF001RDE, ENCFF000HFF, ENCFF000HFH |
| RNA-seq | HCT116 | | ENCODE | ENCFF000DKT, ENCFF000DKW, ENCFF000DKV, ENCFF000DKX, ENCFF000DKY, ENCFF000DKU |
| RNA-seq | HCT116 | | Kelso, 2017 | GSM2719768, GSM2719769 |
| RNA-seq | HCT116 | | Dai, 2018 | GSM2775145, GSM2775146 |
| RNA-seq | Hap1 | | Haarhuis, 2017 | GSM2493886, GSM2493887, GSM2493888, GSM2493898, GSM2493899, GSM2493900 |
| RNA-seq | Hap1 | | Essletzbichler, 2014 | SRX655511, SRX655512 |
